# Visual and auditory object representations in ventral visual cortex after restoring sight in humans

**DOI:** 10.1101/2024.11.22.624459

**Authors:** Katarzyna Rączy, Madita Linke, Job van den Hurk, Carolin Heitmann, Maria J. S. Guerreiro, Minye Zhan, Ramesh Kekunnaya, Rainer Goebel, Brigitte Röder

## Abstract

Visual category-selective representations in human ventral occipital temporal cortex (VOTC) seem to emerge early in infancy. Surprisingly, the VOTC of congenitally blind humans features category-selectivity for auditory and haptic objects. Yet it has been unknown whether VOTC would show category-selective visual responses if sight were restored in congenitally blind humans. Assuming competition for synaptic space during development, cross-modal activation of VOTC as a consequence of congenital blindness might interfere with visual processing in sight-recovery individuals. To test this hypothesis, we investigated adults who had suffered a transient phase of congenital blindness due to bilateral dense cataracts before their sight was restored by cataract-removal surgery. In a functional magnetic resonance imaging (fMRI) study, participants watched movies of faces, scenes, body parts and other objects in the visual condition, while in the auditory condition they listened to the corresponding sounds. The most prominent group difference was the reduced face-selectivity in individuals with reversed congenital cataracts compared to age- and sex-matched normally-sighted individuals. In addition, a double dissociation was found: only sight-recovery individuals demonstrated significant decoding accuracy of visual categories based on auditory category representations in VOTC, while only normally-sighted individuals’ VOTC decoded auditory categories based on visual category representations. The present results uncovered the neural mechanisms of previously observed face processing impairments in individuals with reversed congenital blindness. We suggest that lower face-selectivity in the sight recovery group might arise from selective deficits in the cortical representation of the central visual field in lower-tier visual areas. Additionally, we speculate that in higher-order visual areas cross-modal activity might facilitate – rather than interfere – with visual functional recovery after congenital blindness.

## Introduction

The ventral occipital temporal cortex (VOTC) of the human brain is known to comprise regions selective for different visual object categories. Sub-regions preferentially, though not exclusively, respond to faces (fusiform face area, FFA, (1)), body parts (fusiform body areas, FBA, (2); extrastriate body area, EBA, (3)), scenes (parahippocampal place area, PPA, (4)), and other objects (lateral occipital complex, LOC, (5, 6)). There is currently a debate on whether category-selective representations exist at birth (7) or are a result of domain-independent processing properties that induce category-selectivity in VOTC by visual experience (8). In adult humans (9, 10) and monkeys (11, 12), face-selective regions are retinotopically biased to the fovea; that is, most face-selective neurons have receptive fields near fixation. By contrast, scene-selective regions are dominated by peripheral parts of the visual field (13),(14). These retinotopic biases seem to exist at birth and have been hypothesized to guide the acquisition of category-selective domains within the ventral visual stream during development (8). In fact, if monkeys were deprived of faces after birth, face-selective neural circuits did not emerge (15), suggesting that appropriate experience is necessary for face-selectivity to emerge (13).

In humans, a number of recent studies have demonstrated that infants (2-9 months of age (16)) show some degree of category-selectivity within the VOTC for faces, scenes and body parts, but not for other visual categories such as toys or tools (16, 17). While some studies reported adult-like neural category-selectivity within the first year of life (17–20), others have found that particularly face-selective regions continue to mature until adolescence or even adulthood (7, 21–23). The observation that both the amount and type of face experience affected face processing abilities in human adults (24–26) was counted as evidence for an experience-dependent development of associated neural systems. These results were in line with the proposal that functions that emerge late in development were found to be more susceptible to early visual deprivation than those that emerge early – even if sight was restored before functional development was complete (termed ‘sleeper effect’ by Maurer et al (27)).

Brain imaging studies in congenitally blind humans (e.g. (28–36)) have casted doubts on the assumption that category-selective representations in VOTC are exclusively a result of visual experience and built on retinotopic biases: In the complete absence of visual experience from birth, category-selectivity in VOTC was reported for haptic and auditory object stimuli instead (for review see (36, 37)). The topographic distribution of different categories, moreover, resembled that known for visual objects in normally-sighted individuals (36, 37). For example, van den Hurk et al. (35) were able to decode the category-selectivity for auditory stimuli in individuals with permanent congenital blindness based on the category-selective organization of the VOTC for visual stimuli of normally-sighted individuals and vice versa. These results suggest, on the one hand, that VOTC represents modality-invariant object categories and, on the other hand, that these neural representations are not necessarily dependent on visual input during development.

However, it is yet unknown whether neural representations acquired without visual information are capable of category-selective processing of *visual* objects. Blind humans rely entirely on auditory and haptic input, which might be potent to set up category-selective representations in the VOTC. It was speculated that cross-modal connectivity shapes these category-selective representations (8). Even normally-sighted individuals might integrate visual, auditory and haptic input for shaping category-selective responses in VOTC. It could be hypothesized that, due to competition for synaptic space during development, visual afferents are pruned away if blindness existed at birth. As a result, if sight were restored later in life (see (38, 39)), the visual system would not have the same access to VOTC as in normally-sighted individuals while the latter would still maintain non-visual category-selectivity. Alternatively, cross-modally – that is, auditorily and haptically – induced category-selective regions in the VOTC could comprise a scaffold for visual responses to emerge if sight was restored in congenitally blind humans.

Here we investigated individuals who had been born without pattern vision due to bilateral dense cataracts and who underwent cataract-removal surgery after the 6^th^ month of life. Research in sight-recovery individuals who mostly received surgery not before late childhood reported that cross-modal matching of haptic and visual objects was impaired immediately after sight restoration but emerged with time (40). These behavioral observations suggest that at least some visual experience is necessary to link visual and haptic object knowledge. Held et al. (40), however, did neither test for category knowledge nor for neural representations of visual objects or categories.

Individuals with reversed congenital cataracts are able to recognize objects (41) even if they had been visually deprived for several years prior to sight restoration (42). A recent eye-tracking study in congenital cataract reversal individuals has found overall high object recognition skills, which were better the more typical their visual exploration patterns when freely viewing images of objects (43). The authors argued, as has been suggested for the typical acquisition of object knowledge (44) and by Arcaro & Livingstone (11), that congenital cataract reversal individuals make use of low- and high-level visual feature information while exploring the visual world after sight restoration and that the resulting systematic eye movement patterns allowed them to acquire visual object knowledge. Ossandón et al. (43) did not, however, test category knowledge nor neural representation for visual objects. At the neural level, typical face-selective responses in individuals with reversed congenital cataracts were neither observed with event-related potentials (ERPs (42)) nor in a brain imaging study (45) – and face and object representations did not fully emerge in a single sight-recovery individual who had suffered from blindness starting at the age of 3 years (46, 47). Yet neither visual nor auditory category-selectivity across multiple categories have been investigated in sight-recovery individuals with a history of transient congenital blindness.

Based on the scarce existing literature, we hypothesize that early visual experience is crucial for acquiring well-tuned face-selective neural representations. Category-selective representations might not be needed for object classification but rather might be essential for within-category processing. In fact, face identity processing under conditions when stimulus context and vantage point were manipulated was markedly impaired in individuals with reversed congenital cataracts (48–50). Whether or not category-selective representations would emerge for other object classes after a transient phase of congenital blindness is largely unknown; however, preliminary behavioral evidence suggested that within-category processing is less impaired for other object classes, e.g. for houses (51).

The present functional magnetic resonance imaging (fMRI) study evaluated both visual and auditory object representations in adult individuals who had been born blind due to dense bilateral cataracts and whose cataracts were surgically removed after the age of 6 months but no later than at 48 months. We were thus able to test in the same individuals for a possible cross-modal interference (pruning of visual influence on VOTC due to congenital blindness) vs. cross-modal facilitation (building on auditorily and possibly haptically shaped object representations after sight restoration) in congenitally blind individuals who (akin to a control group of sighted individuals) likewise had extensive periods of visual experience (i.e., following sight restoration). We assessed category-selective representations for four different categories (faces, scenes, body parts and other objects) as participants watched movies in the visual condition or listened to the corresponding sounds in the auditory condition. First, the data were analyzed using a univariate approach (general linear model) to test for the existence of category-selective representations. Second, we adapted a multivariate approach as a highly sensitive measure for detecting neural representations selective for visual categories (52). Third, to assess the similarity of category-selective representations for auditory and visual objects, we examined the degree to which cross-modal decoding of categories was possible in congenital cataract reversal individuals vs. normally-sighted individuals.

For the univariate analysis, we predicted to replicate typical category-selectivity in the VOTC (e.g. (35, 53)) for visual images in normally-sighted individuals but reduced or absent category-selectivity in individuals with reversed congenital cataracts, particularly for faces.

An interference of auditory and visual category-selective representations would be indicated by a lower cross-modal decoding accuracy (that is, auditory-to-visual and visual-to-auditory) in congenital cataract reversal individuals than in normally-sighted individuals. If, however, auditory category-selective representations (54) promote the emergence of visual category-selective representations in congenital cataract reversal individuals, then we would expect a higher auditory-to-visual decoding accuracy in this group.

Category-selectivity was expected for auditory cortex in both groups.

## Materials and Methods

### Participants

We recruited congenital cataract reversal individuals (CC group) and matched normally-sighted individuals (SC group).

The CC group consisted of eight individuals (mean age: 42.0 years, range: 32-48 years, 3 males) with a history of bilateral dense congenital cataracts. All CC individuals had undergone cataracts removal surgery after the age of 6 months (mean age at surgery: 21 months, range: 6-48 months). Mean visual acuity in the better eye at the time of testing, as assessed on the logarithm of the minimum angle of resolution (logMAR) scale in the Freiburg visual acuity and contrast test (FrACT) (55) was 0.65 logMAR (range: 0.16-1.27; the lower the logMAR the better the visual acuity). For detailed participant information, see Table S1. All CC individuals had normal color vision as indicated by no errors made by any of the participants in the administered color classification test (Farnsworth Dichotomous Color Vision Test: Panel D-15 (56)). For the behavioral testing, participants wore their own visual aids; that is, either glasses or contact lenses. During the scanning, if needed, they received MR compatible glasses, which we had ordered for them according to each participant’s prescription.

The eight SC individuals (mean age: 42.0 years, range: 32-48 years, 3 males) were individually matched in age and sex to the CC group. All SC participants had normal or corrected-to-normal visual acuity and normal color vision as tested with Farnsworth Dichotomous Color Vision Test: Panel D-15 (56).

Participants did not report any neurological or chronic diseases and had normal hearing as measured on the day of testing (for one normally-sighted participant we recorded a mild hearing loss in the right ear at 4000 Hz).

All participants provided written informed consent prior to taking part in the study. Participants received monetary compensation for the time of participation, and we covered other expenses such as travel and accommodation costs.

The study was approved by the local ethics committee of the Faculty of Psychology and Human Movement Science, University Hamburg and by the University of Maastricht and was conducted according to the principles laid down in the Declaration of Helsinki (2013).

### Stimuli

The stimuli and procedure are described in detail in (35). Briefly, four stimulus categories – that is, body parts, scenes, faces and other objects – were presented either visually or in the auditory modality in separate runs. Visual and auditory stimuli consisted of 64 short (∼1,800 ms) movie or audio clips, respectively, with 16 unique clips per category. In the body category, for example, movie clips showed a hand scratching a forearm or two hands clapping. In the scene condition, they showed waves crashing on a beach or a restaurant overview. In the face condition, movie clips presented the lower part of the face while laughing or chewing. In the other-objects category, movies depicted a washing machine or a clock. The auditory clips were the same movie clips as used in the visual condition but presented only the audio track (e.g., the sound of clapping hands or “facial” sounds such as laughing or chewing of the visually presented actions). Auditory stimuli were matched in overall sound intensity by normalizing the root mean square of the sound pressure levels. To ensure maximum visibility for CC individuals, the size of the visual stimuli was doubled for both groups, which resulted in 28*17cm onscreen stimulus size. For the full list of video and audio clips we refer to Job van den Hurk, co-author of the current paper.

### Design and Task

Stimulus presentation was controlled with MATLAB R2014a (The MathWorks, Inc., Natick, Massachusetts, United States) and Psychtoolbox-3 (57). Visual stimuli were back-projected onto a translucent screen which was visible to the participants via a mirror attached to the head-coil (Panasonic PT-EZ570, screen resolution: 1920 × 1200 pixels, refresh rate = 60 Hz, viewing distance: 75 cm). Auditory stimuli were delivered via earphones (Sensimetric systems, Sensimetrics corp., Malden). The experimental paradigm was by and large identical to the one implemented by van den Hurk et al. (35). Stimuli were presented in blocks with each block consisting of clips showing stimuli from one of the four categories (faces, scenes, body parts or other objects). Each block consisted of either eight movie or audio clips separated by a 200 ms interstimulus interval. Each run consisted of 4 blocks of each of the four categories, that is, 16 blocks and lasted approximately nine minutes. Between each of the four category blocks a fixation-cross (white, presented on a black background) was presented for 12 seconds. There were four visual and four auditory runs in total: half of the participants started with visual runs and ended with auditory runs while the other half started with auditory runs followed by visual runs. In contrast to van den Hurk et al. (35), two category blocks of the visual runs included chromatic and the remaining two achromatic stimuli. In the following we do not separately analyze chromatic and achromatic visual stimuli to improve the signal to noise ratio. The presentation order was counterbalanced within and between runs to control for possible order effects. During the fMRI scanning, participants performed a one-back task and rated the conceptual dissimilarity of the currently presented stimulus relative to the preceding stimulus on a scale of 1–4 with a corresponding button press (1: very similar, 4: very dissimilar), even across blocks. In the instructions prior to the fMRI session, each participant was given an example: a cat meowing and a dog barking were considered as conceptually similar, and a cat meowing and a car starting were considered as dissimilar. The task aimed at encouraging the participants to attentively process the categorical information of the stimuli. At the start of the scanning session, and after the participants were provided with hearing protection and earphones, the volume level was individually adjusted.

### Data acquisition

Data were acquired at the Scannexus MRI scanning facilities at Maastricht University (Scannexus, Maastricht, The Netherlands) with a 64-channel head-neck coil in a 3-Tesla Siemens Prisma Fit scanner (Erlangen, Germany). Functional data comprised 180 - 217 single-shot echoplanar (EP) images (GRAPPA factor = 2, acquisition matrix = 100*100, FoV = 200*200, voxel resolution = 2 mm isotropic, 64 slices, no inter-slice gap, TR = 2000 ms, TE = 30 ms, multiband factor = 2, flip angle = 77°, activated pre-scan normalization and the phase encoding direction posterior to anterior (P-A)). Before each functional scan, five EP images with equal acquisition parameters but opposite phase encoding direction (i.e. A-P) were acquired for image distortion correction.

Additionally, 3D T1 images were acquired using a MPRAGE sequence (GRAPPA factor = 2, 192 slices, acquisition matrix 256*256, FoV 256*256, iso-voxel resolution 1 mm isotropic, no interslice gap, TR = 2300 ms, TE = 2.98 ms, flip angle = 9°, active pre-scan normalization & (inline) distortion correction).

### FMRI preprocessing and data analysis

#### Preprocessing

The f/MRI data were preprocessed and analyzed using the SPM12 software package. (http://www.fil.ion.ucl.ac.uk/spm/software/spm12/). A standard pre-processing pipeline was employed in which all the acquired functional volumes were realigned using rigid body transformations to correct for head movements. Head movements were <2mm in all but three participants (two SC and one CC participant; z <3.34 mm). None of the participants was excluded due to head movements. The framewise displacement (FD) of the time series did not differ between the groups (SC: M=0.17, SD=0.11; CC: M=0.15, SD=0.08; t(7)=0.55, p=.583). For the univariate analysis (GLM), the data were normalized to the standard adult brain template (MNI space) and smoothed with a 5 mm (FWHM) Gaussian kernel. For the multivariate pattern analysis (MVPA) the normalization and smoothing steps were omitted, and the analysis was performed in native space. For univariate analysis, GLM was run for the visual and auditory categories: body parts, scenes, faces and other objects. The main predictors were convolved with the hemodynamic response function (HRF) and the head movement estimates were entered as confound predictors.

### Category-selectivity

#### Whole-brain analysis

First, we assessed category-selective regions for both the SC and the CC group. Four category-selective contrasts for each group were defined: 1) faces vs. all other categories 2) scenes vs. all other categories, 3) bodies vs. all other categories and 4) other objects vs. all other categories, separately for the visual (Figure 1A) and auditory conditions (Figure S6A-D). Next, to assess whether the category-selective regions significantly overlapped between the CC and the SC group, we performed a conjunction analysis as proposed by Nichols et al. (58): voxels were considered as active if they reached significance level in each of the two groups. These analyses were run for all four visual category-selective contrasts (Figure S1A-D) and four auditory (Figure S6A-D).

**Figure 1.**
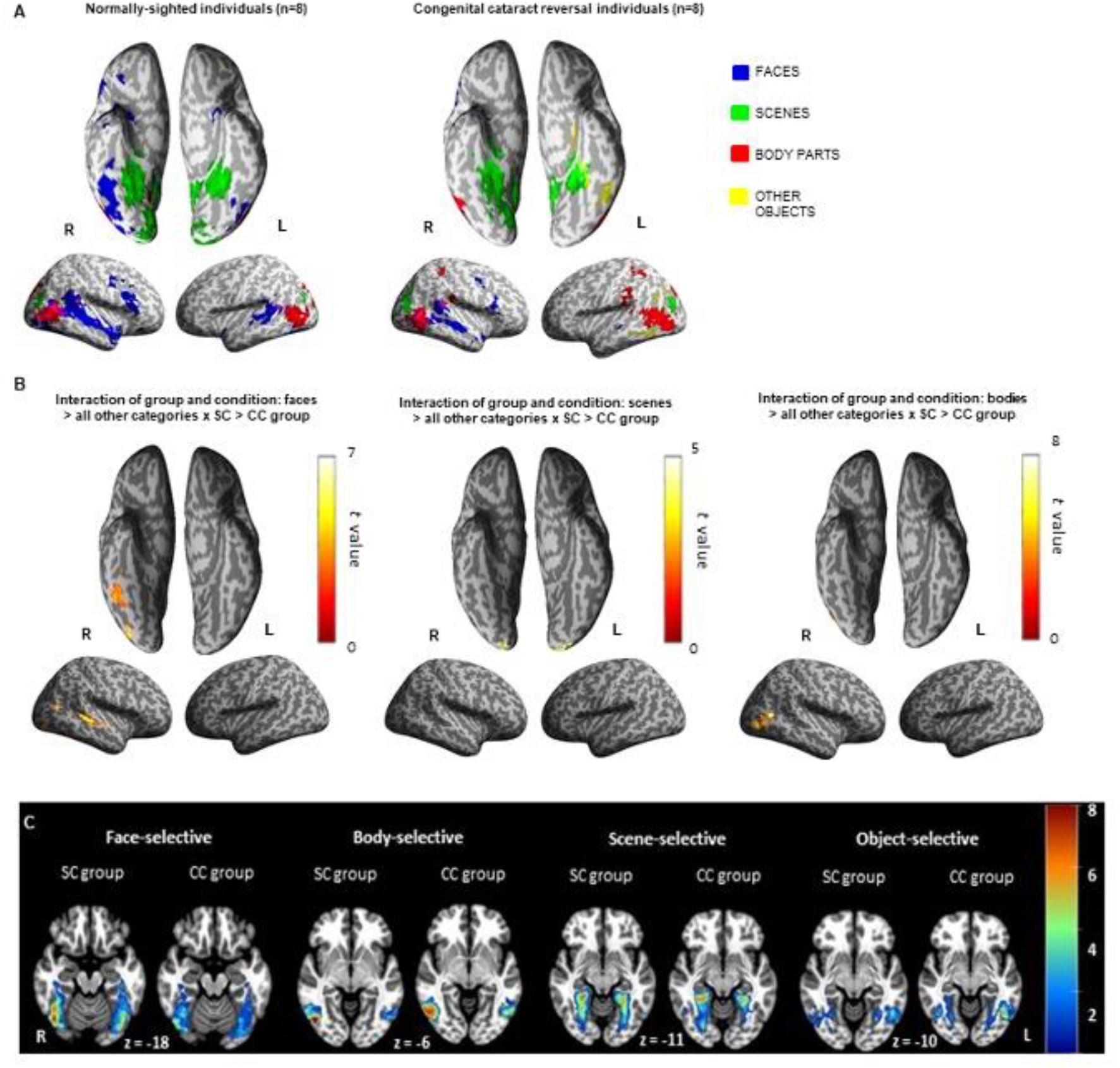
Whole-brain analysis. **A)** Statistical maps for each category-selective contrast were calculated for visually presented movie clips of: faces (vs all other categories), scenes (vs all other categories), bodies (vs all other categories) and other objects (vs all other categories) in normally-sighted individuals (the SC group) and in congenital cataract reversal individuals (the CC group). **B)** Interaction between group and condition. The voxel-wise threshold was set to p < .001 uncorrected and the resulting statistical maps were corrected for multiple comparisons using cluster-wise FWE-correction at p < .05. **C**) Group-overlap maps for face-selectivity, body-selectivity, scene-selectivity and other object-selectivity are shown separately for the SC and the CC group. Individual subject maps of category-selective contrasts were thresholded (p < .0001, voxel-wise, uncorrected) and the resulting statistical maps were corrected for multiple comparisons with cluster-wise FWE-correction at p < .05. Each voxel was consecutively assigned the color which represents the number of the participants who showed an above threshold activation in this voxel. For better visualization, group-overlap maps were constrained by the ventral occipital temporal mask (VOTC; see *Methods: Within-Subject Classification*). L = left hemisphere; R = right hemisphere.

To test for group differences in the category-selectivity effect, we performed a random-effects whole-brain interaction analysis for each visual category with group as factor (category 1 > all other categories x SC group > CC group) (Figure 1B). The whole brain interaction analysis was then repeated for auditory conditions (Figure S6E). The voxel-wise threshold was set to p < .001 uncorrected and the resulting statistical maps were corrected for multiple comparisons using cluster-wise FWE-correction at p < .05.

#### Region of Interest analysis

To compare the response profiles between the CC and the SC group in individually and independently defined cortical regions we applied a region of interest (ROI) approach. First, to independently define ROIs in each participant, the 30 (not necessarily contiguous) most active voxels (that is, with the highest t-value) in each category-selective contrast were identified for each participant in the first visual run. The search for voxels was restricted by the functional mask, as defined in the visual atlas (59) for the corresponding category-selective region. For example, the 30 most active face-selective voxels were identified within the inferior occipital gyrus (IOG), mid-lateral fusiform gyrus (mFus) and posterior-lateral fusiform gyrus (pFus) of each hemisphere as these regions were reported as being face-selective in an independent sample of normally-sighted individuals by Rosenke et al. (59). Given that the visual atlas of Rosenke et al. (59) did not include an object-selective region, in the present study we used the location of the lateral occipital complex (LOC) previously identified by other authors and thus the search for voxels was constrained by a box with coordinates: ymin = -82; ymax = -70; xmin = -45; xmax = 35; zmin = -18; zmax = -9 (following (5, 60, 61). Next, in these voxels we extracted beta activation parameters (z-transformed) from the remaining three visual runs, for each participant and for each experimental condition (i.e. faces, scenes, body parts and other objects). The beta-values were then averaged first across all 30 voxels in each participant and in each condition and then submitted into a mixed-effects ANOVA with group (two levels: congenital cataract reversal individuals, normally-sighted individuals) as a between-group factor and category (four levels: faces, scenes, body parts, other objects) as within-group factor (Figure 2, Table S9-12).

**Figure 2.**
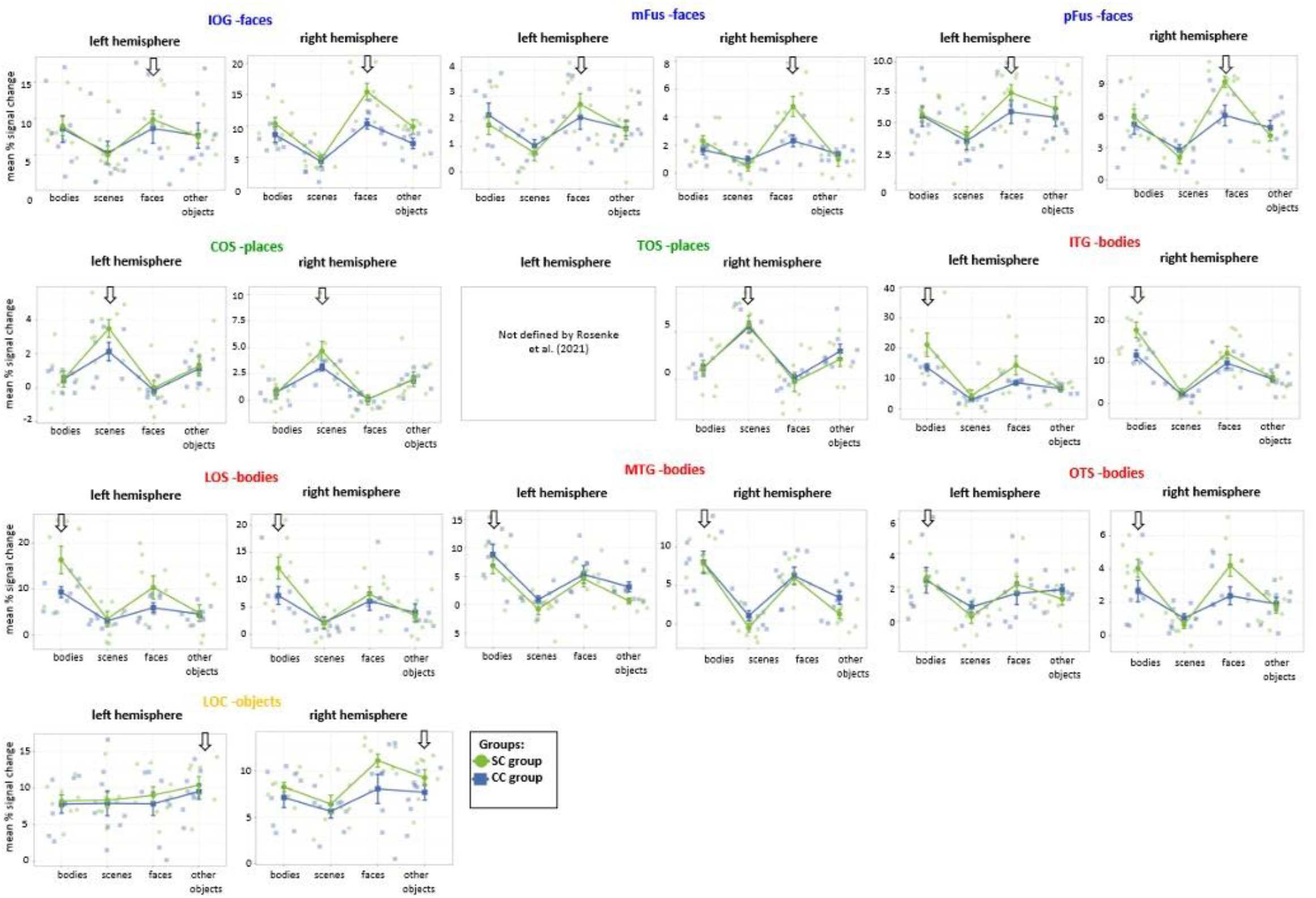
ROI analysis in the regions functionally defined by Rosenke (2021). Mean signal change in category-selective regions is shown for bodies, scenes, faces and other objects in the SC and the CC group. First, to independently define the ROIs the 30 most active voxels in each of the participants were identified within the functional mask, as defined in the visual atlas of Rosenke et al., (2021), in the first visual run. Next, in these voxels we extracted the beta activation parameters from the remaining three visual runs for each participant and each experimental condition. Finally, the beta-values were averaged across all 30 voxels in each participant. CC= congenital cataract reversal individuals, SC = normally-sighted individuals.

Next, to determine whether the reduced face-selectivity of the CC group found in the 30 most active voxels (see above) was consistent across different ROI sizes and in the broader regions of the right fusiform gyrus, we re-ran the analysis with various ROI sizes ranging from 30 to 120 most active voxels (as indicated by the t-values, Figure S5, Table S13-15). Voxels were selected within anatomical borders of the right fusiform gyrus (region FG1, 2, 3, 4). The mask was created with the SPM Anatomy Toolbox version 2.2b (62).

Statistical analyses, as well as visualization of the data, were done in R (version 4.2.1).

#### Multivariate Pattern Analysis

Decoding was performed using a linear support-vector machine (l-SVM) classifier based on the LIBSVM algorithm (https://www.csie.ntu.edu.tw/~cjlin/libsvm/) implemented in The Decoding Toolbox (TDT) version 3.997E (https://sites.google.com/site/tdtdecodingtoolbox) (63).

#### Within-Subject Classification

For the MVPA, we selected four anatomically defined regions of interest (ROI): 1) early visual cortex covering V1 and V2 (areas hOc1 and hOc2), 2) ventral occipital temporal cortex (VOTC; identical as used in (31), 3) primary auditory cortex including superior temporal gyrus (regions TE 1.0, 1.1, 1.2) and 4) higher auditory cortex including superior temporal gyrus (region TE 3). Anatomical masks 1, 3 and 4 were created with the SPM Anatomy Toolbox version 2.2b (62) while VOTC mask (2) was retrieved from available data (31). The decoding was performed with feature selection: A filter method was implemented which used classification weights on features (here referred to as voxels) in a training set to determine the contribution of each voxel in the given classification. Next, the number of high performing voxels in each training set was individually estimated and these voxels were then used in the classification, both in the training and the testing set. These voxels were then used in the classification. Decoding was performed in a pairwise manner: first for all possible pairs and then averaged over all four categories. If during the classification there was a tie (i.e. no assignment to any category) between two categories in a pairwise comparison, the undecided test case was added to the separate column of the confusion matrix. We used a leave-one-run-out cross-validation: the decoding was run four times, such that each run was used once as test data and the remaining three runs as training data. The final classification accuracy was computed by averaging over the four iterations.

Two-sample t-tests were used to test the significance of differences between the groups in the classification accuracy and one-sample t-tests to test the classification accuracies against the chance level (25%).

#### Cross-Modal Classification

To determine whether cross-modal classification was possible in the CC and in the SC group, the classifier was trained with the data of visual runs and tested on the data of auditory runs and vice versa in each group. The procedure was identical to the within-subject classification with the exception that now all visual/auditory runs were used as training data.

#### Data Quality Assessment

We assessed the quality of fMRI scans (e.g. (64)) to rule out that observed group effects could be due to different data quality: data quality was assessed as the temporal signal-to-noise ratio (tSNR) during the visual runs applying the left and right hemisphere VOTC mask. TSNR did not significantly differ between groups (SC: M=139.99, SD=35.20; CC: M=138.72, SD=20.64; t(14)=0.08, p=.931).

#### Behavior in the scanner

To verify that participants processed stimulus categories we estimated the mean dissimilarity ratings for two conditions, either when a new category block had just begun (between category) or for ratings recorded within a category block. When multiple button presses were registered after a trial, only the first response was counted. Numerically, dissimilarity ratings of both groups were higher when a new block had started than within a block, both in the visual and in the auditory condition. Due to a high drop-out rate, we only descriptively report the rating results of participants who gave at least five out of sixty between-block responses, which included the data of two CC and six SC participants in the visual condition, and the data of six CC and eight SC participants in the auditory condition (Figure S10).

## Results

### Category-selectivity for visual objects: Whole-brain analysis

First, we identified brain regions that selectively responded to each of the four visual categories – that is, faces, bodies, scenes and other objects – in both normally-sighted individuals (the SC group) and congenital cataract reversal individuals (the CC group) (Figure 1A).

In the SC group, face-selective regions were found bilaterally in the occipito-temporal cortex; that is, in the fusiform gyrus and inferior occipital gyrus and in the superior and middle temporal gyrus (Figure 1A, Table S2). Body-selective regions were observed bilaterally in lateral occipito-temporal cortex and in the right parietal cortex (Figure 1A, Table S2). For the scene-selective contrast, several clusters located medially along the inferior occipito-temporal cortex, in the early visual cortex, and parietal regions of both hemispheres were identified (Figure 1A, Table S2). For the other object-selective contrast, no significant cluster in the VOTC was obtained (Figure 1A, Table S2; but see uncorrected category-selective maps in Figure S3A). In the CC group, the univariate analysis did not reveal face-selective regions in the VOTC (Figure 1A, Figure S3B) and thus no overlap between the groups existed here (Figure S1C). Outside the VOTC, face-selective regions were found in middle temporal gyrus and in middle frontal cortex in this group (Figure 1A, Table S2). The former region overlapped between the CC and the SC group (Figure S1C, Table S2). Body-selective regions were found bilaterally in the lateral occipito-temporal and parietal cortex (Figure 1A, for a list of individual brain regions see Table S2) and overlapped between groups (Figure S1A). The scene-selective contrast in CC individuals resulted in significant clusters located bilaterally and medially along the inferior occipito-temporal cortex and in early visual regions of the right hemisphere (Figure 1A, Table S2), which overlapped with those of the SC group (Figure S1B, Table S2). The CC group showed significant clusters in the other object-selective contrast in a region covering left middle occipital gyrus and two regions that in SC individuals selectively responded to faces, that is, left fusiform gyrus, and left inferior temporal gyrus (Figure 1A, Table S2). No overlapping clusters for the other object-selective contrast existed between groups (Figure S1D, Table S2).

To directly test for group differences in category-selective processing, a random-effects whole-brain interaction analysis was conducted for each visual category with group as factor (category x > all other categories × SC > CC group). We observed significant group differences in face-selectivity in four clusters of the right hemisphere: in the inferior temporal gyrus and the inferior occipital gyrus, face-selective neural responses were absent in the CC group as compared to the SC group (Figure 1B, Figure S2A, Table S2). While all matched SC individuals (n = 8) showed above-threshold face-selective activation in typical face-selective regions of the right fusiform cortex, only four of the CC individuals did (see Figure 1C). The superior and middle temporal gyrus featured a reduced face-selectivity in the CC group as compared to the SC group (Figure 1B, Table S2). Moreover, the magnitude of selective neural response was greater to bodies and scenes in the SC group as compared to the CC group in the right middle temporal gyrus for bodies (Figure 1B, Figure S2C, Table S2) and in the middle occipital gyrus and left inferior occipital gyrus for scenes (Figure 1B, Figure S2B, Table S2). No significant interaction effects were found for other objects (Figure 1C, Figure S1E).

In sum, the CC group, in contrast to the SC group, did not feature face-selectivity in VOTC (Figure 1A-B) and showed reduced scene- and body-selective responses outside of VOTC (Figure 1A-B). No regions with higher neural selectivity for any of the categories were identified in the CC compared to the SC group.

### Category-selectivity for visual objects: Region of interest (ROI) analysis

Considerable anatomical and functional inter-individual variability have previously been reported, in particular for FFA (e.g. (65)). In order to exclude the possibility that such sources of variability caused the observed interaction effects in the whole-brain analyses, we compared the response profiles of the 30 most active voxels between the SC and the CC groups. Following previous studies (66, 67) the most active 30 voxels were defined as those with the largest t-value for each participant and for each category-selective contrast within functionally constrained category-selective regions as defined by the functional atlas of Rosenke et al. (59): These authors identified category-selective regions by contrasting each category with the mean activation to all other categories in their group of 19 healthy individuals (for details see (59)). Rosenke et al. (59) revealed face-selective regions within mid-lateral fusiform gyrus (mFus) and posterior-lateral fusiform gyrus (pFus), which overlapped with the fusiform face area (FFA (7)), and in the inferior occipital gyrus (IOG), which corresponded to the occipital face area (OFA). The same authors (59) identified place-selective regions in the collateral sulcus (COS), which overlapped with the parahippocampal place area (PPA (4)), and in the transverse occipital sulcus (TOS (68)). Body-selective regions of Rosenke et al., (for the full list see (59)) corresponded to the fusiform body area (FBA (2, 69)) and the extrastriate body area (EBA (3)). Since in the study of Rosenke et al. (59) no sufficient selective activation was found for the other object condition, here we used the location of the lateral occipital complex (LOC) previously suggested to be object-selective by other authors (5, 60, 70). In the present ROI analysis, we considered a region to be category-selective if the average activation of voxels within the predefined region (those of Rosenke et al. (59) and for the other object category defined by (5, 60, 70)) was significantly greater for one (called the preferred) category than for any of the other categories (called the non-preferred categories).

#### Face-selective regions

In the SC group we found larger activation for faces, relative to bodies, scenes and other objects in mFus, pFus and IOG of the right hemisphere (Table S11). This face-selective response was only found in the SC group (significant group x condition interaction; Table S9) but not in the CC (Table S11). As a consequence, the magnitude of cortical responses to faces was larger in the SC than in the CC group in these regions (Table S12, Figure 2).

#### Place-selective regions

Both the SC and the CC group featured scene-selectivity in the COS and TOS of the right hemisphere and the SC group in addition in the left COS (Table S11, Figure 2).

#### Body-selective regions

Body-selectivity was found bilaterally in the inferior temporal gyrus (ITG), middle temporal gyrus (MTG) and lateral occipital sulcus (LOS) of both groups (Table S11). No body-selectivity was found bilaterally in the occipital temporal sulcus (OTS) either in the SC or the CC group (Table S11, Figure 2).

#### Object-selective regions

The SC and the CC group showed significantly larger activation to other objects than to scenes but not to faces or bodies in the right LOC and larger activation to other objects than to bodies but not to faces or scenes in the left LOC, and thus no other object-selectivity was found in these regions in either of the two groups (Table S11, Figure 2).

We did not find any group difference in the magnitude of cortical responses to the non-preferred stimulus categories for neither of the category-selective ROIs analyzed (Table S12).

In an additional analysis the size of the ROIs in the search for face-selective voxels was varied between 30 and 120 voxels (see Figure S5 and Supp Table 13-15) and we extended the search for these voxels to the whole right fusiform gyrus. The group x condition interaction emerged regardless of the ROI size and extent, confirming the robustness of the group differences reported here (Supp Table S13).

In sum, the SC group featured face-selectivity in the previously identified face-selective ROIs of the right hemisphere (as defined by (59)); that is, faces elicited significantly larger activation than any other category. By contrast, in the CC group face-selectivity was not significant in these regions, confirming the whole-brain results. Scene-selectivity was observed in both groups in typical place-selective ROIs (as defined by (59)), though in fewer ROIs in the CC group. Both the SC and the CC group showed body-selectivity in most of the previously identified body-selective regions. We did not find other-object selectivity in either group.

### Category-selectivity for auditory objects: Whole brain analysis

In this paragraph, face-, body-, scene- and other object-selectivity allude to the representations of the corresponding auditory stimuli.

In the SC group, face-selective regions were found bilaterally in the middle and superior temporal gyrus, middle frontal gyrus and inferior parietal cortex (Figure S6C, Table S3). Neither the body-selective nor the scene-selective contrast revealed any significant cluster in the SC group (see Figure S6A-B, Figure S4A). Other object-selective responses emerged mostly in the middle and inferior temporal gyrus and inferior occipital gyrus of the left hemisphere (Figure S6D, Figure S4A, Table S3). CC individuals demonstrated face-selectivity bilaterally in the middle and superior temporal gyrus as well as in the frontal cortex, body-selectivity bilaterally in temporal and frontal cortices, and scene-selectivity in the left fusiform gyrus and the precuneus (Figure S6A-C, Figure S4B, Table S3). Other objects-selectivity was not observed in this group either (Figure S6D, Figure S4B, Table S3). The conjunction analysis revealed an overlap in face-selective responses bilaterally in the superior temporal gyrus and in the left temporal pole (Figure S6C, Table S3).

A subsequent whole-brain interaction analysis found a higher selectivity for faces in the SC group than in the CC group in the superior temporal gyrus of the right hemisphere (Figure S6E, Table S3). Group differences were not found for scenes, bodies or other objects.

### Within-Subject Classification for visual and auditory objects

The multivariate pattern analysis (MVPA) revealed significant classification of all visual categories for both the SC and the CC group in all predefined regions of interest (early visual cortex, ventral occipital temporal cortex (VOTC), primary and secondary auditory cortex; for the full description of ROIs see *Methods*, Figure 3A, Table S5). Classification accuracy in VOTC was marginally higher in the SC group than in the CC group (Figure 3A, Table S5). Confusion matrices confirmed that while both groups featured above chance classification accuracies for each visual category within the VOTC, classification accuracy in VOTC was significantly lower for faces in the CC than in the SC group (Figure S7A, and Table S6). In the early visual cortex, classification accuracy for all visual categories was above chance in both groups; other objects were significantly better classified in this region by the SC than by the CC group (Figure S7A, Table S6).

**Figure 3.**
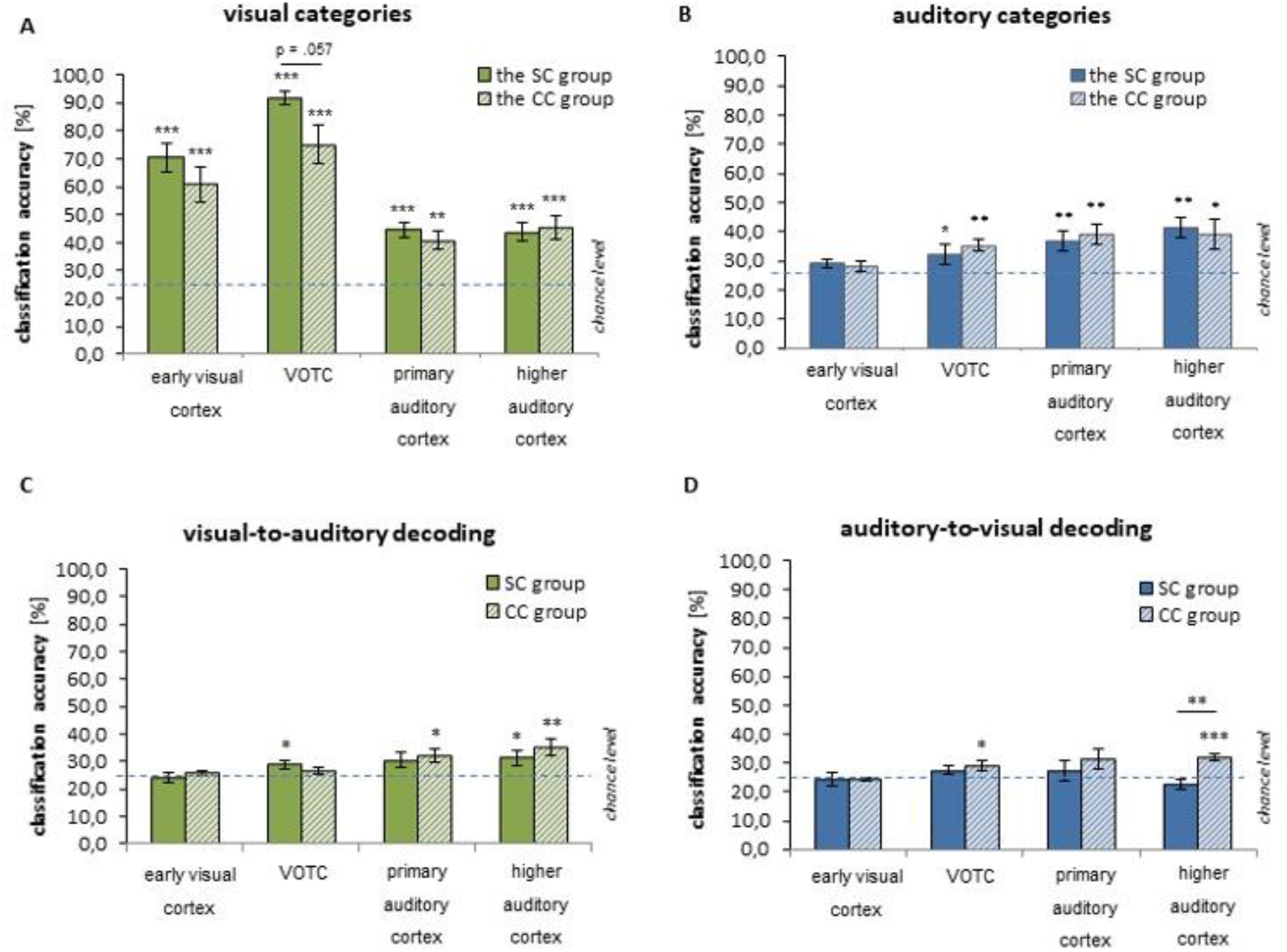
Classification results. Within-subject decoding results for visual A) and auditory B) categories in four ROIs in the CC and the SC group. Cross-modal classification results for C) visual-to-auditory D) and auditory-to-visual decoding in four ROIs in the CC and the SC group. The values were tested against chance level = 25% and between the groups. *p < .05, **p < .01, ***p < .001. undec = undecided cases. CC = congenital cataract reversal individuals. SC = normally-sighted individuals.

All auditory categories were successfully classified in the SC and the CC group by VOTC and the primary and higher auditory cortex (Figure 3B, Figure S7B, Table S5-6) but not by the early visual cortex. Classification accuracies were indistinguishable between groups in all ROIs.

Whole-brain searchlight analysis (for details see Supp Materials) confirmed for both groups that clusters with high decoding accuracy for visual categories were most predominantly located within visual and parietal regions (Figure S8A, Table S4), while clusters with high decoding accuracy for auditory categories were mostly found in auditory cortices (Figure S8B, Table S4).

### Cross-Modal Classification

Decoding of auditory categories based on visual category representations (visual-to-auditory cross-modal classification) was possible in higher auditory cortex for both the SC and the CC group (Figure 3C, Figure S9A, Table S7-8). In VOTC visual-to-auditory classification was significant only for the SC group. By contrast, visual-to-auditory classification was significant for the primary auditory cortex only in the CC group. Decoding of visual categories based on auditory category representations (auditory-to-visual cross-modal classification) was only successful in the CC group in VOTC and in higher auditory cortex and was significantly better in higher auditory cortex in the CC group than in the SC group (Figure 3D, Figure S9B, Table S7-8).

## Discussion

The present study tested whether category-selectivity for visual objects as typically found in the ventral occipital temporal cortex (VOTC) depends on early visual experience and whether cross-modal auditory category-selectivity interferes with or facilitates the recovery of visual category-selectivity in VOTC after restoring sight in congenitally blind humans. Sight recovery individuals with a history of congenital cataracts and a matched group of normally-sighted individuals watched or listened to movie clips representing four different stimulus categories: faces, scenes, body parts and other objects.

Functional magnetic resonance imaging (fMRI) revealed category-selectivity for scenes and body parts in congenital cataract reversal individuals (CC group), which although somewhat reduced, was found in regions similar to those of the normally-sighted control group (SC group). By contrast, for faces we found a particularly strong reduction of selectivity within typical face-selective regions of the VOTC, including fusiform gyrus in the CC group. With multivariate pattern analysis (MVPA) we were able to successfully decode all visual categories in the VOTC of both groups. Nevertheless, in accord with the univariate analysis, decoding of faces was less accurate in the CC group. Finally, auditory categories were successfully classified by both the auditory cortex and VOTC in both groups, with no significant differences between CC and SC individuals. However, successful cross-modal decoding, i.e., decoding of visual categories in VOTC and auditory cortex using a classifier trained on auditory stimuli, was observed only in the CC group. Conversely, cross-modal decoding of auditory categories in VOTC was only significant in the SC group.

### Category-selectivity is predominantly reduced for faces in the VOTC of CC individuals

Previous developmental fMRI studies in both humans (71) and monkeys (72) have suggested that category-selectivity in the VOTC typically predominantly arises from a decreased responsiveness to non-preferred stimulus categories rather than an enhanced responsiveness to the preferred category. Notably, face processing skills in children improved as a function of the loss of responsiveness of the right fusiform gyrus to non-facial stimuli (71). In accord with this idea, neurophysiological (42) and brain imaging (45) results in individuals with reversed congenital cataracts found that typical brain circuits for faces similarly processed facial and non-facial stimuli. The N170 amplitude, considered as a marker of the structural encoding of faces, showed similar amplitudes for faces, houses and scrambled images in CC individuals and thus was higher for non-facial stimuli in this group compared to normally-sighted individuals (42). These results gave rise to the hypothesis that early visual experience is critical for the tuning of face-selective regions. However, neither this study nor the fMRI study of Grady et al. (45) were designed to investigate genuine category-selectivity in VOTC for faces and other stimulus categories.

A decrease in responsiveness to non-preferred stimuli (71) was considered to involve structural changes such as synaptic pruning following a phase of overproduction of synapses during development (73). Synaptic pruning, but not proliferation, was demonstrated to be experience-dependent in non-human primates (74). The higher cortical thickness observed in the visual cortex of permanently blind humans, which has consistently been reported in previous studies (e.g. (75–78)), suggests a lack of experience-dependent pruning and remodeling. Recent research has proposed that increased myelination may account for developmental changes in cortical thickness (79), including in regions associated with face and word processing (80). Sight-recovery individuals have been found to exhibit higher cortical thickness, similar to congenitally blind individuals (39, 76). Thus, the lower functional selectivity for faces in the VOTC of CC individuals may partially arise from an impaired structural remodeling. This would indicate that sensitive periods for structural and functional tuning of cortex go hand in hand.

Recent research (80) has indicated that the functional remodeling likely includes an enhancement of the response to the preferred category as well. Thus, the reduced category-selectivity to faces might be due to a reduced structural remodeling and as a consequence, a lack of functional tuning for faces.

In the current study, category-selective response was found for body parts and scenes in CC individuals. Although selectivity for these two categories was somewhat reduced, the group difference was far less pronounced than for faces. First, in the whole brain analysis CC individuals featured body parts- and scene-selectivity but no face-selectivity in any sub-region of the VOTC. Second, decoding accuracy for faces but not for body parts and scenes was significantly reduced in the CC compared to the SC group indicating less distinct distributed neural representation in CC individuals (81).

Here we speculate why category-selectivity for faces is particularly dependent on early visual experience. A potential explanation could be related to altered retinotopic biases which are known to exist in different category-selective regions. In normally-sighted individuals, population receptive field (pRF) sizes in ventral face-selective regions have been shown to be densely packed at the center of the visual field; that is, they seem to be mostly driven by foveal visual input (10). While an adult retinotopic organization in early visual areas has been found to be in place in early childhood (at the age of 5 years (10, 13)), the foveal bias for faces in the fusiform gyrus seems to emerge later in childhood (10). Thus, if the foveal but not the more peripheral aspects of visual field representations were altered in CC individuals, particularly face-selectivity would be predicted to be reduced in VOTC. In fact, our recent high-field fMRI study (82) has demonstrated a selective impairment of foveal representations in the very same CC individuals who participated in the present study: pRF sizes were larger in the CC group for the foveal region and, unlike in the SC individuals, pRF sizes did not increase towards the parafoveal visual field. Moreover, the cortical magnification factor for the foveal region was over proportionally reduced in the CC group (82). Interestingly, Heitmann et al. (82) did not observe the typical increase in pRF size towards downstream areas in CC individuals, which they interpreted as a lack of pooling of visual information along the visual processing hierarchy. An earlier EEG study in an independent cohort of sight recovery individuals came to the same conclusion (83). Pooling of information from lower tier visual areas, however, is crucial for face identity processing (84, 85). Although we did not specifically assess face identity processing, previous research in CC individuals (50, 86), one study including a few participants of the present study (50), demonstrated intact face detection but impaired face individuation in CC individuals ((50), see (86) too). A pooling of high-resolution information arising from the fovea seems to be important for the extraction of face-invariant features and for constructing a holistic representation of faces (49).

Therefore, we speculate that the altered representation of the central visual field and degraded pooling across retinal regions contributed to the reduced face-selectivity in face-processing regions of the VOTC (such as the fusiform gyrus) in the CC group. As discussed in Heitmann et al. (82), lower pRF sizes for the central visual field cannot be explained by retinal abnormalities but rather originate at the cortical level. Moreover, largely preserved category-selective representations for body parts and scenes are in accord with the indistinguishable pRF sizes for parafoveal regions of CC and SC individuals. In fact, pRF sizes decreased rather than increased for foveal to parafoveal visual regions in the CC group, excluding visual acuity deficits or the effects of nystagmus, which are typically observed in CC individuals, as accounts for the present findings and those of Heitmann et al ((82); see there for a more detailed discussion).

Interestingly, we observed selective responses for other objects in CC individuals in regions of the left hemisphere, which partially corresponded to typical face-processing areas in normally-sighted individuals. This result is reminiscent of an observation by Grady et al. (45) and aligns with a previous observation in face-deprived monkeys: in face-deprived monkeys, typical face-selective regions responded to other objects long after the face deprivation had ended. As discussed above, longitudinal studies in human children reported that regions of the fusiform gyrus were equally responsive to faces and objects (87) until an adult-like face-selectivity emerged in late childhood. It could be hypothesized that faces and other visual categories compete for synaptic space within the VOTC during development. In the absence of e.g. face or any type of visual object stimulation, the fusiform gyrus might maintain responses to other visual categories. The situation resembles observations in illiterates: not only that the Visual Word Form Area did not emerge in this group, but the fusiform gyrus retained a higher responsiveness to other visual categories as well (88, 89).

### Preserved face-processing outside the VOTC of CC individuals

Face-selective regions were found in the superior temporal sulcus (STS) of the CC group, although to a lower degree as compared to the SC group. This finding echoes an earlier fMRI study, which did not find lip-reading specific responses in the STS of CC individuals (50). Interestingly, Grady et al. (45) observed an altered functional connectivity between the fusiform face area and other regions of face-processing network in CC individuals. Thus, it seems plausible to speculate that the reduced face-selectivity in the fusiform gyrus of CC individuals might be related to the lower selectivity of superior temporal sulcus for face actions such as reading lip movements. Lateral face-selective regions have been shown to rely to a larger degree on more peripheral than foveal input; that is, the pRF profile of the superior temporal sulcus is more similar to that of scene-selective regions than of ventral face-selective regions (9). Since Heitmann et al. (82) reported preserved pRF sizes for parafoveal space in CC individuals, it might be speculated that the reduced though still significant face selectivity in the CC group’s superior temporal sulcus was associated with the higher reliance on non-foveal visual input.

Classification accuracy for faces in the early visual cortices did not differ between the SC and the CC group. This outcome is in line with results from congenitally face-deprived monkeys for which V1 activity was reported to look normal after the end of the deprivation (15). These findings align well with two recent EEG studies in CC individuals which provided evidence for a higher recovery of striate than extrastriate cortical processing (83, 90). In the present context, indistinguishable classification accuracies for faces in early visual areas in CC and SC individuals highlight the genuine processing impairment in VOTC and undoubtedly rule out potential confounding factors related to visual acuity or similar factors.

### Cross-modal decoding of visual categories in the VOTC of CC individuals

Category-selectivity in the visual cortex for auditory and haptic stimuli was reported for congenitally permanently blind humans (31, 35–37) and, in fact, was shown to largely overlap with visual category-selective regions of normally-sighted individuals (35, 91).

In our study, we found that auditory categories were discriminated by the VOTC of both SC and CC individuals. Interestingly, cross-modal decoding of visual categories by auditory representations was only possible in the CC group’s VOTC and by their higher auditory cortex. These findings are reminiscent of an earlier behavioral observation of Guerreiro et al. (92), who assessed auditory and visual motion aftereffects after adapting to a visual and auditory motion stimulus, respectively. A cross-modal visual motion aftereffect was only found in CC individuals. The authors interpreted this result as evidence for stronger cross-modal auditory-to-visual cortex connectivity due to a transient period of congenital blindness. Similarly, Bruns at al.

(93) has recently reported that CC individuals not only use vision to calibrate auditory spatial representations but, additionally, recalibrated visual spatial representations based on auditory spatial information. Here we speculate that auditory object representations may facilitate rather than interfere with visual category-selective representations in VOTC. Similarly, the SC group featured cross-modal decoding from visual to auditory category information. This result replicated, the often-observed visual dominance in object processing (94). Thus, we can conclude that the typical visual dominance in object processing did not emerge in CC individuals. The cross-modal decoding analyses, thus, revealed a double dissociation: better auditory-to-visual decoding in CC individuals but better visual-to-auditory decoding in the SC group. This double dissociation provides the strongest possible evidence for the specific effects of early sensory experience: visual input results in a visual dominance in object processing while in case of a lack of visual input after birth, auditory input takes over and promotes later visual processing in case of sight restoration. The double dissociation, moreover, excludes the involvement of unspecific factors for the observed group differences such as lower visual acuity.

### Limitations

Sight recovery individuals deprived from visual input from birth for more than 6 months are an extremely rare population of subjects, particularly in Western countries. Results based on a small sample size of eight participants should thus be considered as preliminary. However, there are a number of reasons that speak in favor of valid results. First, in the SC group we were able to replicate the typical category-selectivity for all used categories in regions reported in previous studies. Second, in the CC group we observed selectivity for body parts and scenes in regions overlapping with those for the SC group. Third, lower face-selectivity in VOTC of CC individuals is in accord with preliminary electrophysiological (42) and brain imaging studies (45). Fourth and finally, we observed successful cross-modal decoding of the visual categories based on the auditory representations in the VOTC of CC individuals and successful cross-modal decoding of auditory categories based on visual representation in the VOTC of SC individuals, corroborating the robustness of the current results.

## CONCLUSION

We observed a diminished face-selectivity in sight-recovery individuals with a history of congenital blindness while category-selectivity for body parts and scenes was more preserved. As a reason we discuss an impaired cortical representation of the central visual field and a lack of integration of foveal input in downstream areas. Crucially, we observed enhanced cross-modal decoding of visual categories in individuals with reversed congenital cataracts, suggestive of a supportive rather than interfering influence of cross-modal activity in higher-order visual cortex after restoring sight in congenitally blind humans.

## Acknowledgments

We thank Cordula Hölig for valuable discussions about the data analysis and the results. We thank the department of ophthalmology of the Medical Center Hamburg-Eppendorf, in particular Frank Schüttauf, Diana Bittersohl, and Hanna Faber, for participant recruitment.

