## Supplementary Information for "Visual and auditory object representations in ventral visual cortex after restoring sight in humans"

### Supplementary Figures

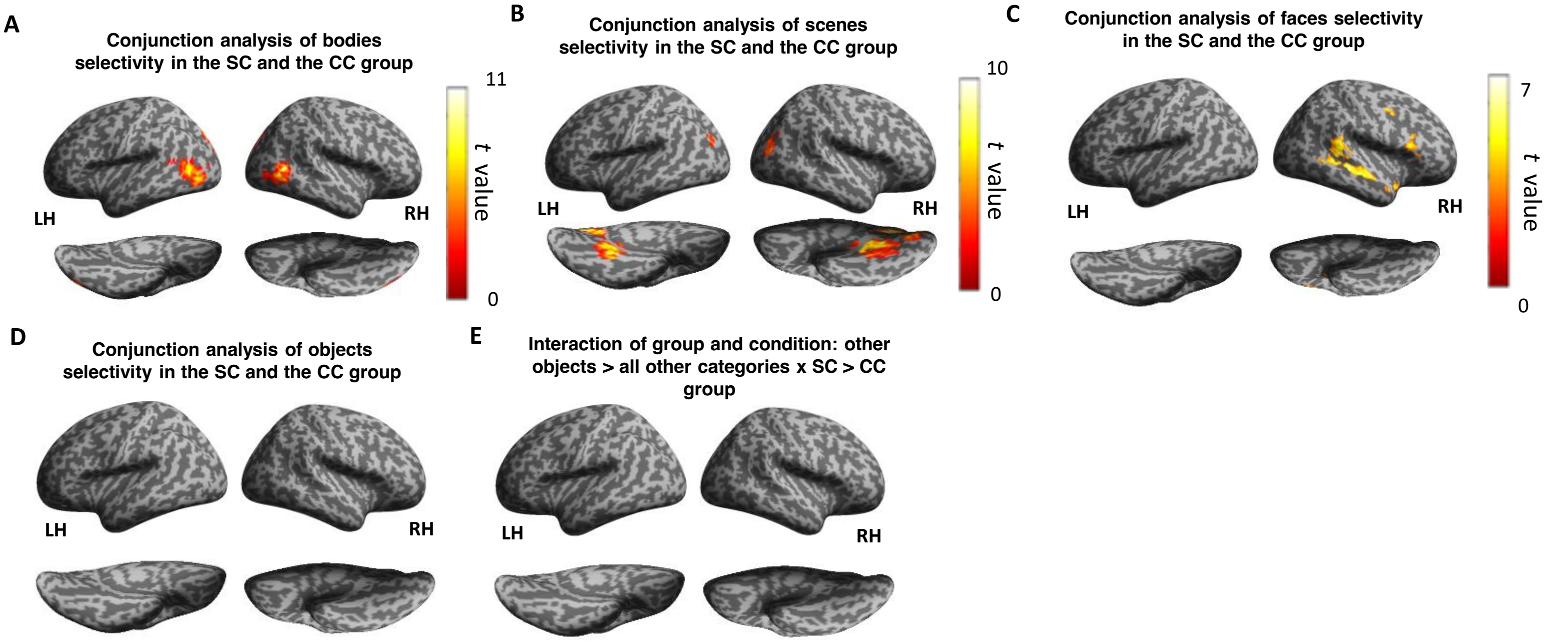


**Figure S1. Whole brain analysis.** Conjunction analysis presents the brain regions activated in the category-selective contrast for: **A**) bodies vs all other categories, **B**) scenes vs all other categories, **C**) faces vs all other categories and **D**) other objects vs all other categories, in normally-sighted individuals (the SC group, n=8) and in congenital cataract reversal individuals (the CC group, n=8). **E**) Interaction between group and condition. The voxel-wise threshold was set to $p < .001$ uncorrected and the resulting statistical maps were corrected for multiple comparisons with cluster-wise FWE-correction at $p < .05$. LH = left hemisphere; RH = right hemisphere.

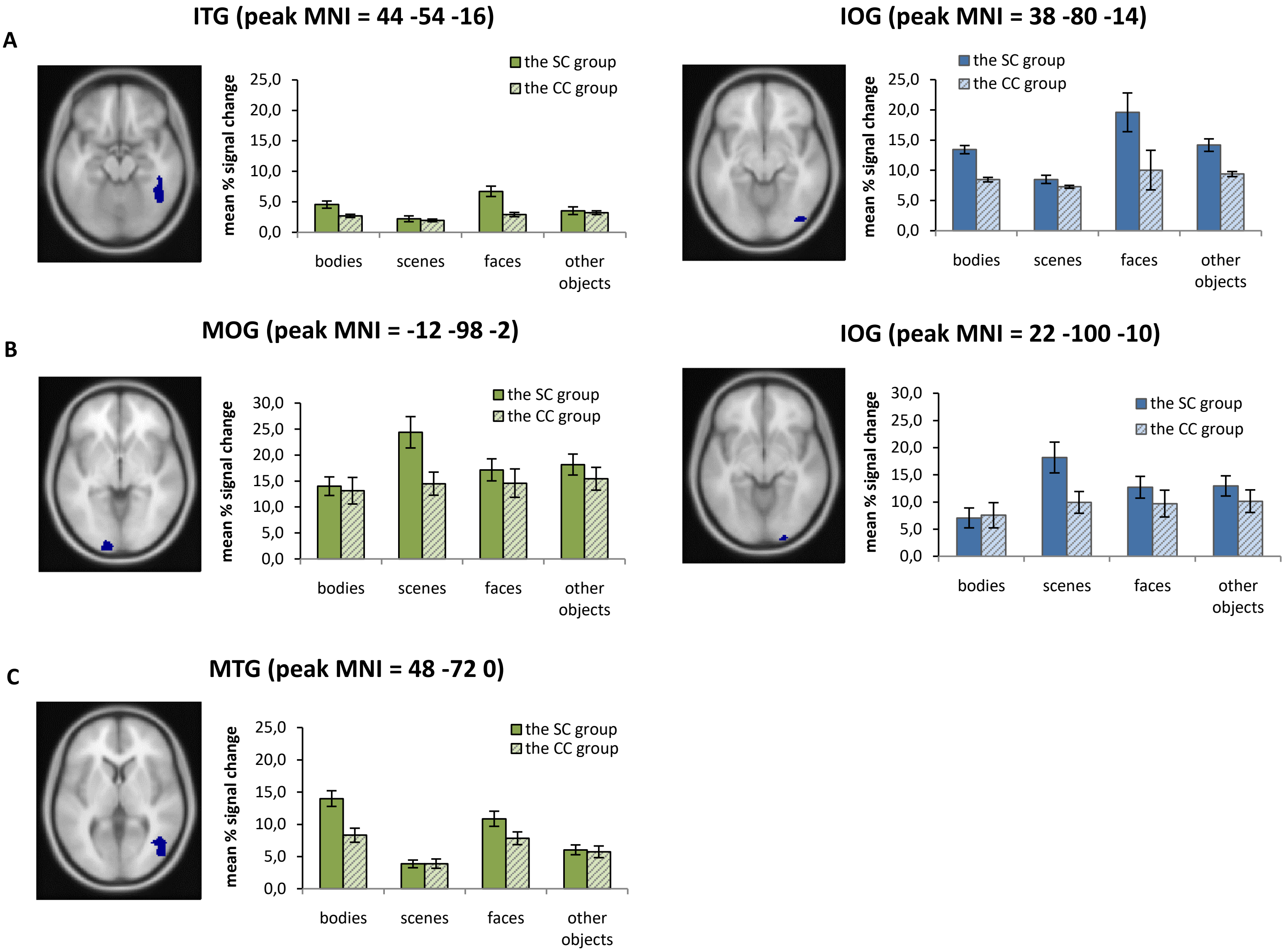


**Figure S2. Mean normalized activation values.** Normalized (Z-transformed) beta-values were first averaged across voxels within the clusters showing significant interaction effects of group and condition in whole-brain analysis, and next across participants in each group and are presented for each visual category. Clusters with the visual cortex are shown for A) face-selectivity, B) scene-selectivity and C) body-selectivity. Error bars represent SEM. ITG = Inferior temporal Gyrus, IOG = Inferior occipital Gyrus, MOG = Middle Occipital Gyrus, MTG = Middle Temporal Gyrus, CC= congenital cataract reversal individuals, SC = normally-sighted individuals.

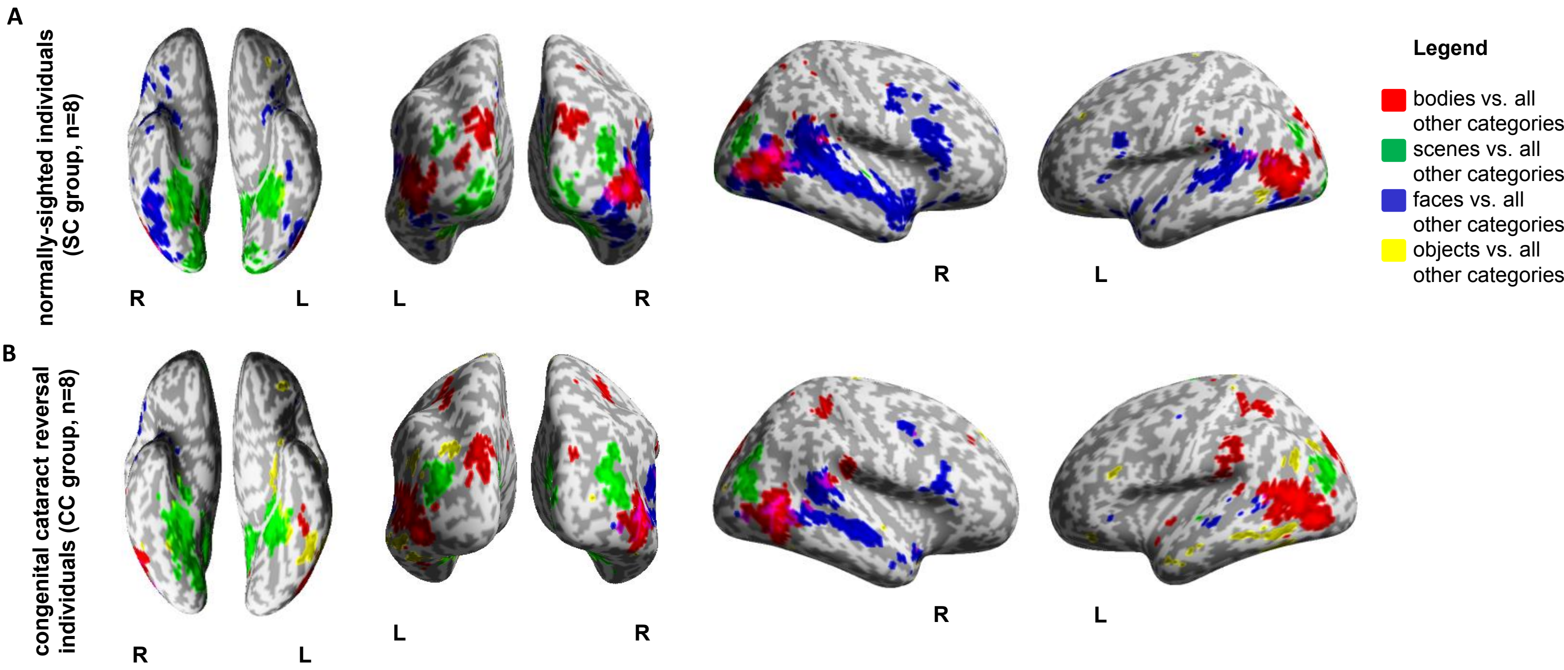


**Figure S3. Category-selectivity maps for visually presented categories** in **A**) the SC group (normally-sighted control individuals) and **B**) the CC group (congenital cataract reversal individuals). The category-selectivity maps visualize the functional topography of the brain for the four categories (bodies, scenes, faces and other objects) presented visually to the participants in each of the tested groups. These maps were created for visualization purpose only. Threshold: $p < .001$, uncorrected. R = right hemisphere; L = left hemisphere.

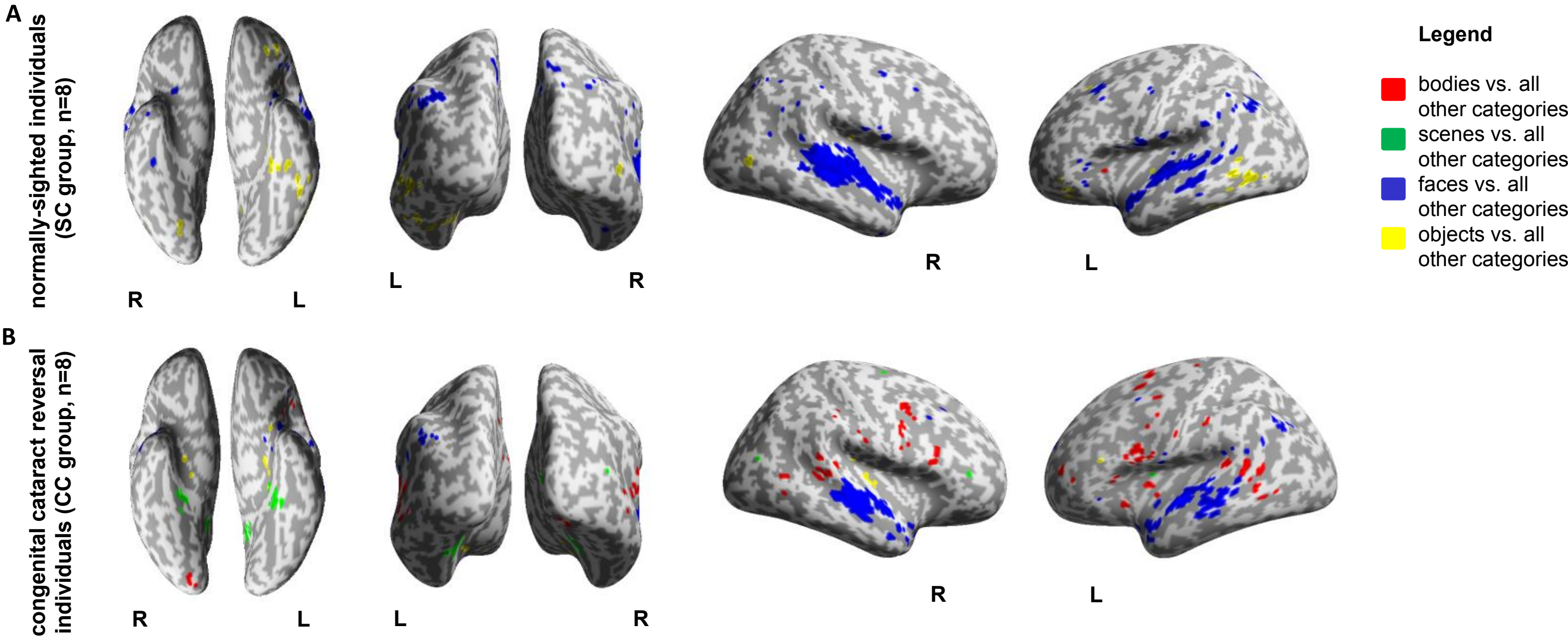


**Figure S4. Category-selectivity maps for auditorily presented categories** in **A**) the SC group (normally-sighted control individuals) and **B**) the CC group (congenital cataract reversal individuals). The category-selectivity maps visualize the functional topography of the brain for the four categories (bodies, scenes, faces and other objects) presented visually to the participants in each of the tested groups. These maps were created for visualization purpose only. Threshold: $p < .001$, uncorrected. R = right hemisphere; L = left hemisphere.

**A**

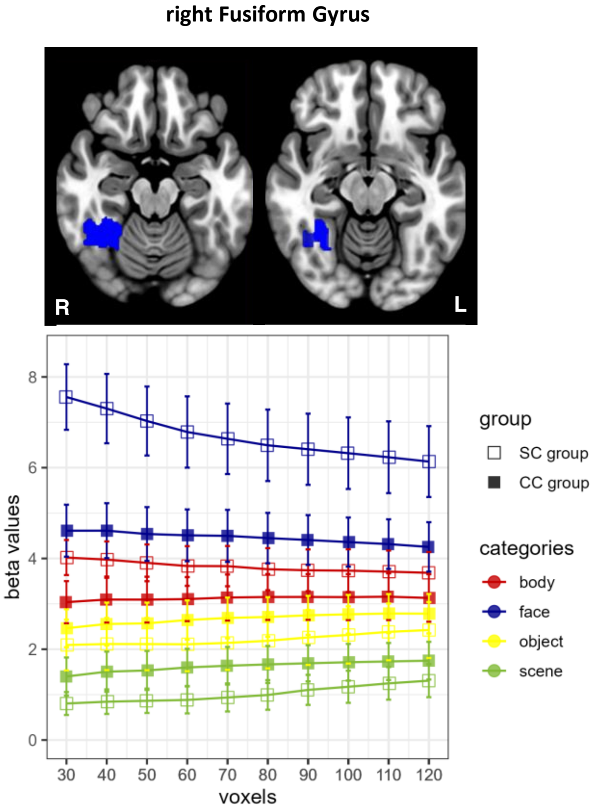


**Figure S5. ROI analysis of the right Fusiform Gyrus for various ROI sizes.** Mean signal change of the face-selective voxels, identified within the anatomical boarder of the right Fusiform Gyrus, is shown for bodies, scenes, faces and other objects in the SC and the CC group in various ROI sizes ranging from 30 to 120 most active (non-contiguous) voxels. The search for these voxels was extended to the whole right fusiform gyrus to ensure that highly face-selective voxels were not omitted due to the constrains of the masks defined by Rosenke et al. (2021). CC = congenital cataract reversal individuals, SC = normally-sighted individuals.

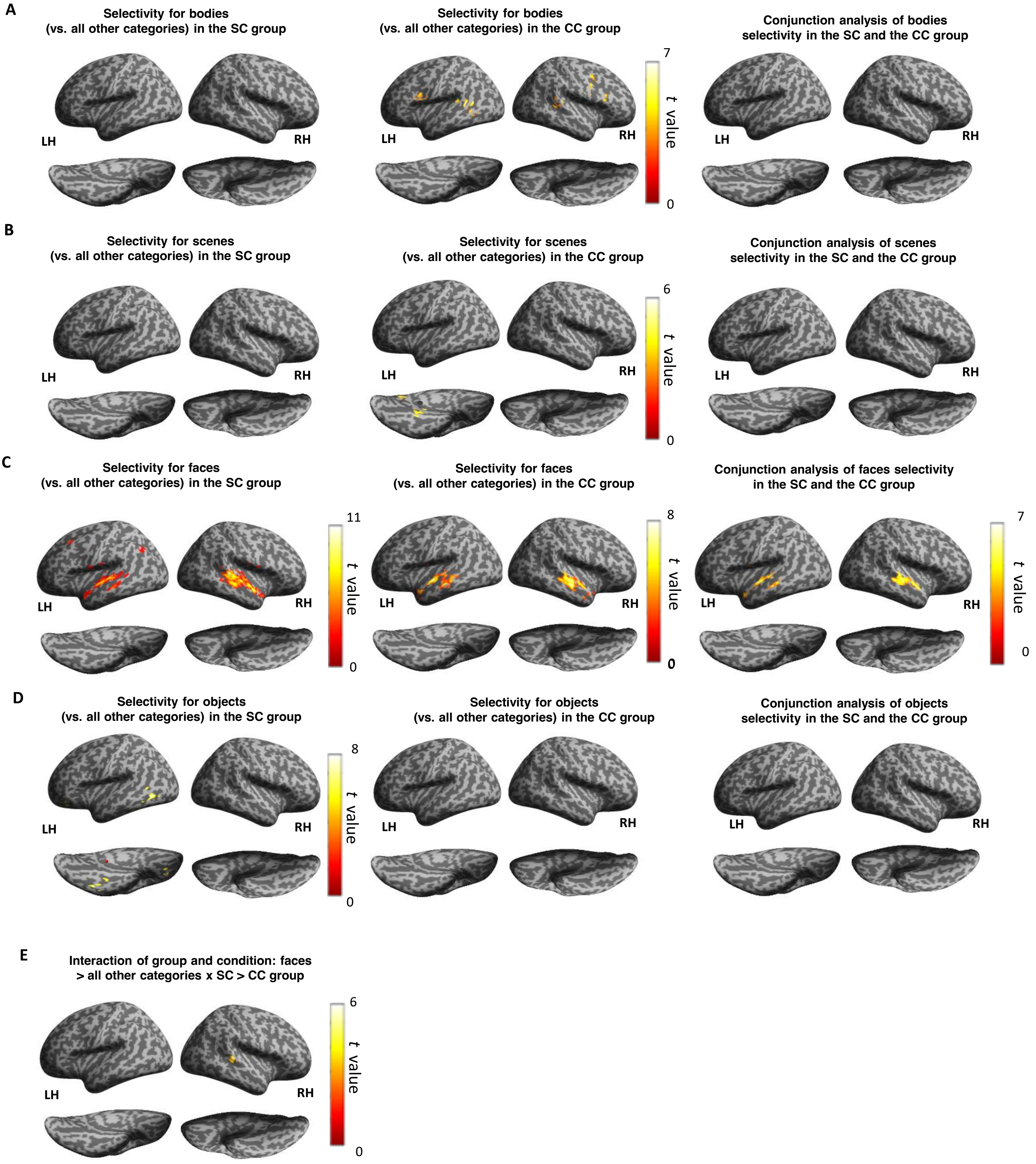


**Figure S6. Whole brain analysis.** Category-selectivity maps were calculated for auditorilly presented movie clips: **A**) bodies vs all other categories, **B**) scenes vs all other categories, **C**) faces vs all other categories and **D**) other objects vs all other categories, in normally-sighted individuals (the SC group, n=8) and in congenital cataract-reversal individuals (the CC group, n=8). Conjunction analysis (third column) presents the brain regions activated in the category-selective contrast in both groups. **E**) Interaction between group and condition. The voxel-wise threshold was set to $p < .001$ uncorrected and the resulting statistical maps were corrected for multiple comparisons using cluster-wise FWE-correction at $p < .05$. LH = left hemisphere; RH = right hemisphere.

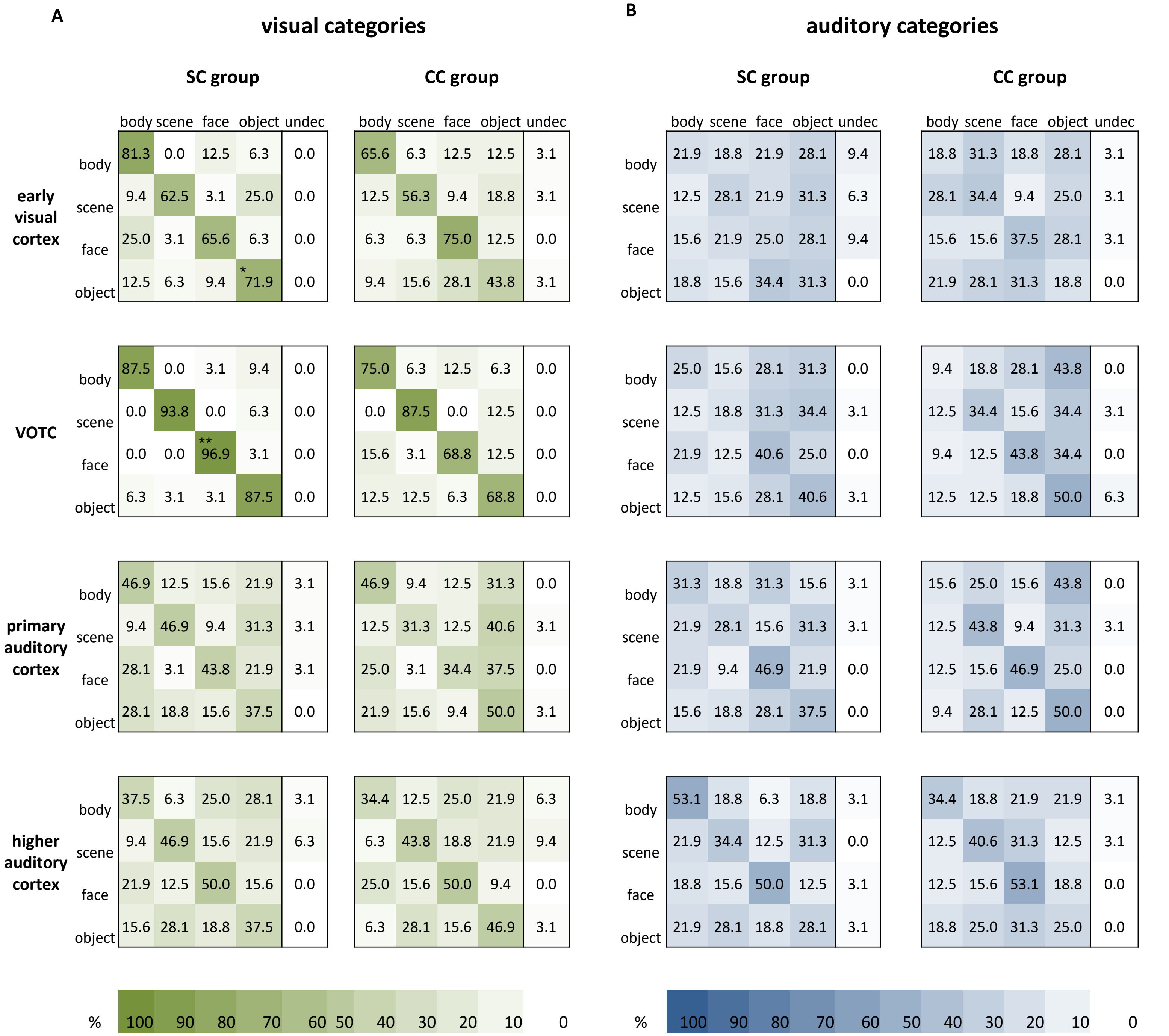


**Figure S7. Within-subject classification results.** Unbiased confusion matrices which exclude undecided pairs in the classification for **A**) visual and **B**) auditory categories in the SC (normally-sighted individuals) and the CC (congenital cataract reversal individuals) group. The vertical row indicates the true category and the horizontal row the predicted category. The axial row represents the accuracy for each category vs all other categories. The values were tested against chance level = 25% (significance not indicated in the confusion matrices, see corresponding Table S6) and between the groups. *p < .05, **p < .01, ***p < .001. undec = undecided cases.

**Searchlight Analysis**

A multivariate whole-brain searchlight analysis was performed to investigate the distribution of voxels with the highest decoding accuracy for the within-subjects classification between four categories (i.e. bodies, scenes, faces and other objects) in visual and auditory condition. The decoding was done in a native space and separately for each participant. A feature selection with a filter method was implemented: classification weights on features (i.e. voxels) in a training set were used to determine the contribution of each voxel in the given classification. Next, the number of high performing voxels in each set was individually estimated. These voxels were then used in the classification. The searchlight was defined as a sphere with a 4-mm radius which moved sequentially across the brain until it was centered at each voxel in the brain. A leave-one-out cross-validation (i.e. the decoding was run four times, each run was used once as test data and the remaining three runs as training data) was performed at each position of the searchlight and the classification accuracy was stored in the center voxel of the searchlight. Then the classification was repeated for the next voxel. As a result, classification accuracy for each voxel located in the center of the sphere was obtained. These whole-brain accuracy maps for each participant were then subsequently normalized to the standard adult brain template (MNI space) and smoothed with a 5-mm (FWHM) Gaussian kernel. The resulting participants' accuracy maps, were then analyzed at the group level, adopting a standard random-effects approach. The analysis was performed separately for visual and auditory condition.

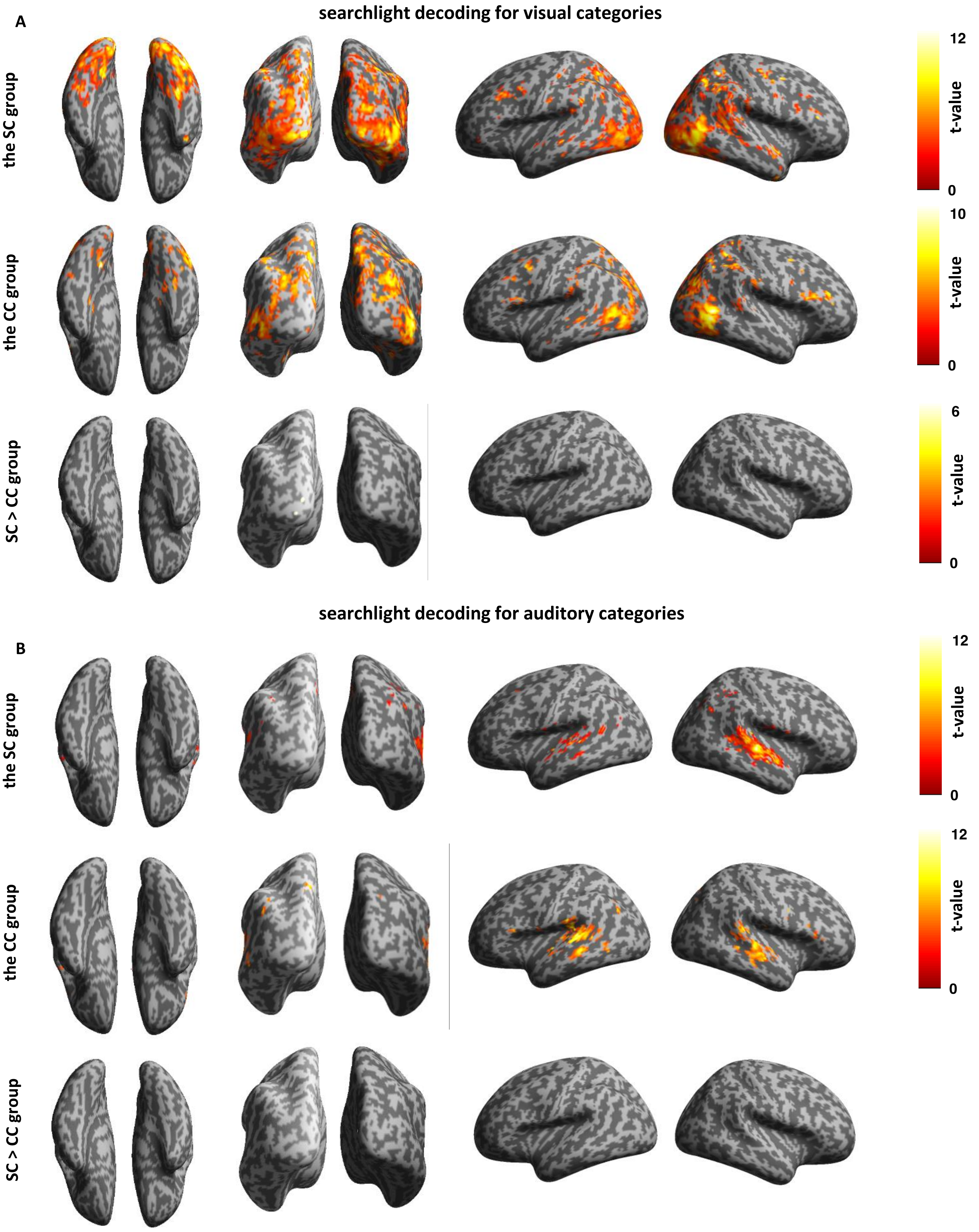


**Figure S8. Whole-brain searchlight analysis.** Results for A) visual and B) auditory categories in normally-sighted individuals (SC group) and in congenital cataract reversal individuals (CC group). Clusters with high decoding accuracy for visual categories were most predominantly located within visual and parietal regions while clusters with high decoding accuracy for auditory categories were found in auditory cortices in both groups. Clusters with high classifier performance are shown at $p < .0001$ uncorrected voxel wise and at $p < .05$ FWE-corrected cluster wise.

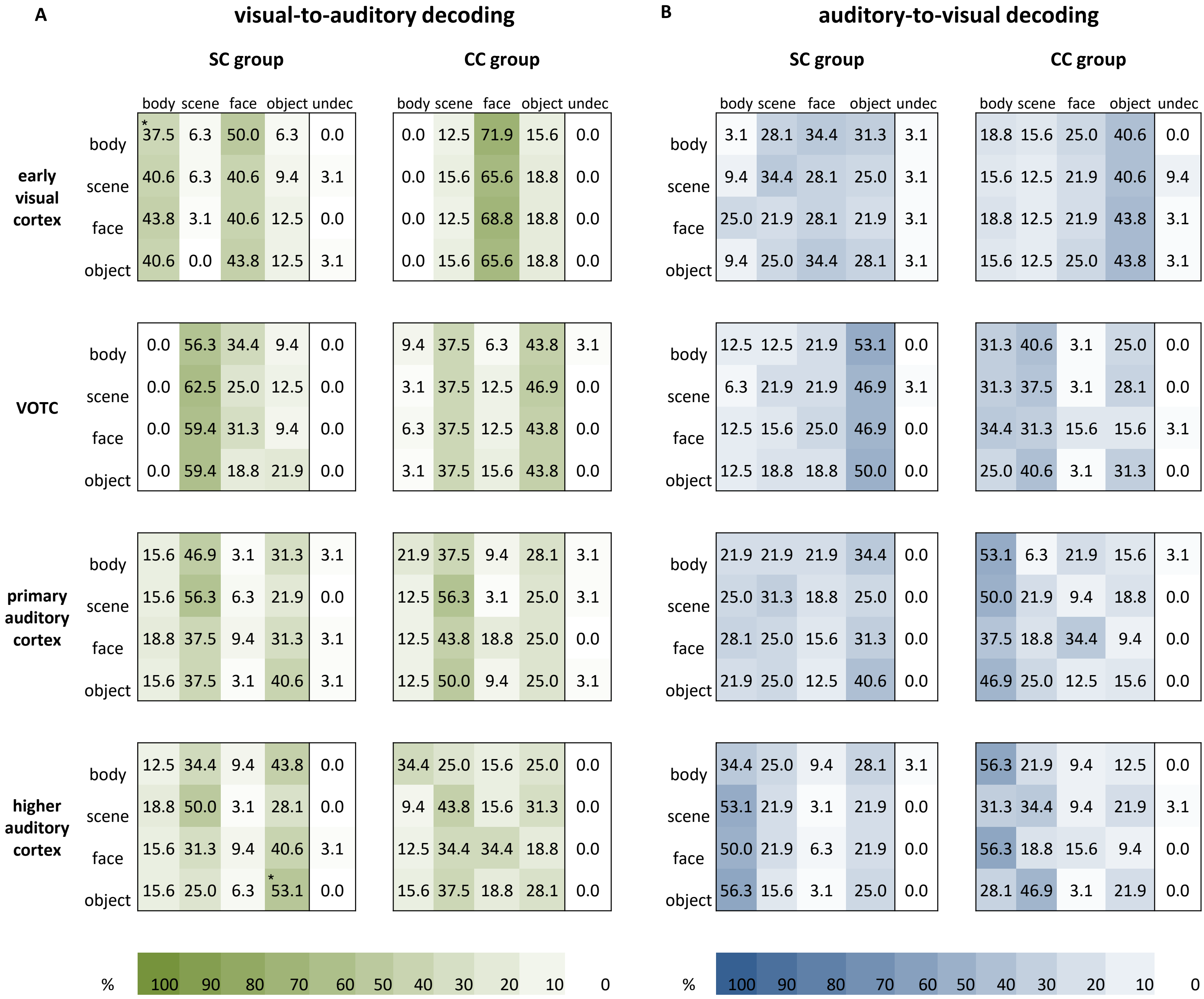


**Figure S9. Cross-modal classification.** Unbiased confusion matrices that exclude undecided pairs in the classification for **A**) visual-to-auditory and **B**) auditory-to-visual classification in the SC (normally-sighted individuals) and the CC (congenital cataract reversal individuals) group. The vertical row indicates a true category and the horizontal row the predicted category. The axial row represents the accuracy for each category vs all other categories. The values were tested against chance level = 25% (significance not indicated in the confusion matrices, see corresponding Table S8) and between the groups. *p < .05, **p < .01, ***p < .001. undec = undecided cases.

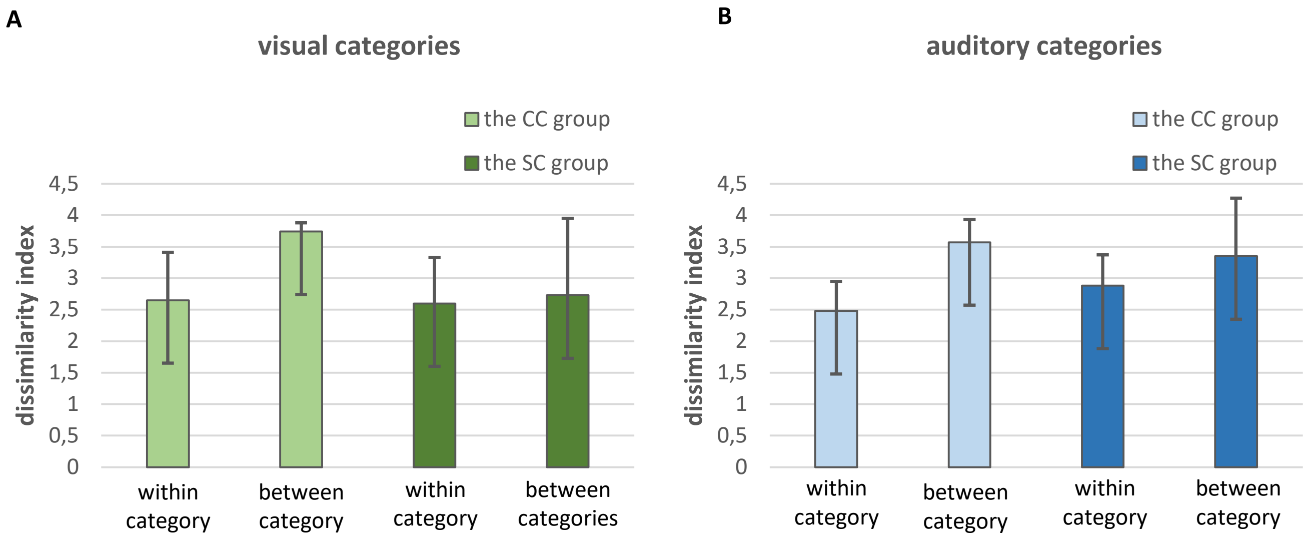


**Figure S10. Behavior in the scanner.** To verify that participants perceived the stimuli between categories as more dissimilar as within categories, a dissimilarity index was estimated: After each stimulus participants rated its dissimilarity to the previous stimulus on a scale from 1 – very similar to 4 – very dissimilar. The mean dissimilarity rating is shown in (A) for visual categories and in (B) for auditory categories. Numerically, dissimilarity ratings of both groups were higher when a new block had started than within a block, both in the visual (A) and in the auditory (B) condition. Error bars represent SEM. SC = normally-sighted individuals, CC = congenital cataract reversal individuals.

### Supplementary Tables

**Table S1**. Clinical and demographic characteristics of congenital cataract reversal individuals.

| ID | Sex | Age at testing (years) | Age at surgery (month) | Visual Acuity (decimal) | Visual Acuity (logMAR) | Additional details |
|---|---|---|---|---|---|---|
| CC1 | M | 34 | 12 | 0.24 | 0.63 | Aphakia |
| CC2 | M | 45 | 24 | 0.70 | 0.16 | Aphakia, impaired stereo vision |
| CC3 | F | 48 | 48 | 0.26 | 0.58 | Aphakia, impaired stereo vision |
| CC4 | F | 45 | 9 | 0.21 | 0.68 | Pseudoaphakia, nystagmus, treated glaucoma |
| CC5 | F | 32 | 24 | 0.66 | 0.18 | Aphakia |
| CC6 | F | 46 | 24 | 0.05 | 1.27 | Aphakia, nystagmus, impaired stereo vision, microphthalmia |
| CC7 | F | 46 | 18 | 0.07 | 1.16 | Aphakia, nystagmus, impaired stereo vision, monocular enucleation |
| CC8 | M | 40 | 6 | 0.28 | 0.56 | Aphakia, nystagmus, impaired stereo vision |

*Note.* M = male; F = female, logMAR = logarithm of the minimum angle of resolution.

**Table S2.** Brain regions with significant clusters in category-selective contrasts in the SC and the CC group for the visual runs.

| Contrast | Hemisphere | Brain Region | Brodmann Area | Cluster size | t Statistics | MNI coordinates | | |
|---|---|---|---|---|---|---|---|---|
| | | | | | | x | y | z |
| **Bodies vs. other categories in the SC group** | | | | | | | | |
| | R | Middle Temporal Gyrus | 37 | 1317 | 15.2 | 48 | -60 | 4 |
| | | | | | 13.68 | 50 | -72 | -2 |
| | | | | | 10.33 | 42 | -64 | 0 |
| | L | Middle Occipital Gyrus | 19 | 1260 | 10.18 | -46 | -76 | 6 |
| | | | | | 8.83 | -42 | -84 | 2 |
| | | | | | 4.15 | -58 | -58 | 12 |
| | L | Superior Occipital Gyrus | 19 | 752 | 9.15 | -16 | -88 | 34 |
| | L | Middle Occipital Gyrus | 19 | | 4.86 | -20 | -86 | 16 |
| | L | Cuneus | 19 | | 4.47 | -10 | -86 | 22 |
| | R | Superior Occipital Gyrus | 19 | 491 | 7.69 | 22 | -84 | 34 |
| | | | 7 | | 7.30 | 26 | -76 | 34 |
| | R | Cuneus | 19 | | 5.09 | 12 | -74 | 34 |
| | R | Superior Temporal Gyrus | 22 | 124 | 5.23 | 62 | -38 | 16 |
| | R | SupraMarginal Gyrus | 39 | | 4.27 | 58 | -48 | 28 |
| | | | 40 | | 3.65 | 54 | -42 | 24 |
| | R | Lingual Gyrus | 18 | 87 | 4.76 | 12 | -60 | -4 |
| | R | Calcarine Gyrus | 17 | | 3.89 | 12 | -66 | 8 |
| | R | Cuneus | 18 | | 3.50 | 12 | -70 | 20 |
| **Bodies vs. other categories in the CC group** | | | | | | | | |
| | L | Inferior Occipital Gyrus | 19 | 1586 | 9.37 | -44 | -72 | -4 |
| | L | Middle Temporal Gyrus | 19 | | 8.87 | -46 | -64 | 4 |
| | L | Middle Occipital Gyrus | 19 | | 8.35 | -50 | -74 | 6 |
| | R | Middle Temporal Gyrus | 37 | 1080 | 11.02 | 54 | -64 | 2 |
| | | | | | 10.46 | 50 | -58 | -4 |
| | R | Inferior Temporal Gyrus | 37 | | 6.99 | 60 | -58 | -8 |
| | L | Superior Occipital Gyrus | 19 | 558 | 6.67 | -22 | -84 | 38 |
| | | | | | 6.41 | -20 | -88 | 30 |
| | L | Cuneus | 23 | | 4.77 | -12 | -80 | 32 |
| | L | Superior Temporal Gyrus | 22 | 437 | 5.32 | -60 | -38 | 14 |
| | | | 40 | | 5.22 | -54 | -32 | 20 |
| | L | SupraMarginal Gyrus | 40 | | 4.89 | -56 | -28 | 36 |
| | R | Superior Temporal Gyrus | 40 | 383 | 6.04 | 62 | -28 | 20 |
| | R | Rolandic Operculum | 40 | | 5.43 | 52 | -30 | 22 |
| | R | Superior Temporal Gyrus | 22 | | 4.76 | 64 | -36 | 16 |
| | L | Inferior Parietal Lobule | 7 | 303 | 5.87 | -28 | -46 | 50 |
| | L | Superior Parietal Lobule | 7 | | 5.43 | -38 | -48 | 60 |
| | L | Postcentral Gyrus | 1 | | 5.07 | -30 | -36 | 52 |
| | R | Postcentral Gyrus | 1 | 173 | 6.83 | 26 | -36 | 46 |
| | | | | | 3.32 | 32 | -38 | 60 |
| | R | Superior Occipital Gyrus | 19 | 88 | 4.60 | 20 | -80 | 32 |
| **Conjunction analysis: body selectivity in the SC group AND in the CC group** | | | | | | | | |
| | L | Middle Occipital Gyrus | 19 | 986 | 8.35 | -50 | -74 | 6 |
| | L | Middle Temporal Gyrus | 19 | | 7.66 | -48 | -66 | 6 |
| | L | Middle Occipital Gyrus | 19 | | 7.23 | -48 | -74 | -2 |
| | R | Middle Temporal Gyrus | 37 | 658 | 10.28 | 52 | -62 | 2 |

| | | | | | | |
|---|---|---|---|---|---|---|
| | 19 | | 6.65 | 46 | -68 | 2 |
| | | | 5.98 | 44 | -60 | -2 |
| L Superior Occipital Gyrus | 19 | 376 | 6.41 | -20 | -88 | 30 |
| L Cuneus | 19 | | 5.79 | -18 | -80 | 36 |
| **Group comparison: bodies > other categories x SC > CC group** | | | | | | |
| R Middle Temporal Gyrus | 19 | 352 | 7.37 | 48 | -72 | 0 |
| | | | 6.28 | 48 | -62 | 6 |
| | | | 4.74 | 42 | -64 | 0 |
| **Scenes vs. other categories in the SC group** | | | | | | |
| R Calcarine Gyrus | 17 | 3995 | 10.38 | 16 | -54 | 10 |
| R Lingual Gyrus | 18 | | 9.97 | 28 | -46 | -8 |
| R Calcarine Gyrus | 17 | | 9.22 | 10 | -86 | 0 |
| L Precuneus | 18 | 1557 | 11.9 | -16 | -58 | 14 |
| L Fusiform Gyrus | 37 | | 8.53 | -20 | -44 | -12 |
| L Lingual Gyrus | 18 | | 8.06 | -26 | -52 | -8 |
| R Middle Occipital Gyrus | 19 | 314 | 5.74 | 44 | -76 | 22 |
| | | | 5.62 | 38 | -70 | 22 |
| | | | 5.60 | 40 | -82 | 16 |
| L Middle Occipital Gyrus | 19 | 221 | 5.85 | -34 | -80 | 24 |
| **Scenes vs. other categories in the CC group** | | | | | | |
| R Precuneus | 23 | 2235 | 12.2 | 18 | -54 | 18 |
| L Fusiform Gyrus | 37 | | 11.99 | -24 | -38 | -16 |
| R Precuneus | 23 | | 10.71 | 12 | -54 | 10 |
| R Fusiform Gyrus | 37 | 1759 | 11.04 | 28 | -32 | -20 |
| R Calcarine Gyrus | 17 | | 9.74 | 8 | -84 | 0 |
| R ParaHippocampal Gyrus | 36 | | 8.67 | 20 | -36 | -12 |
| R Middle Occipital Gyrus | 19 | 572 | 7.99 | 40 | -78 | 22 |
| | 39 | | 7.06 | 42 | -76 | 32 |
| | | | 4.58 | 34 | -82 | 38 |
| L Middle Occipital Gyrus | 19 | 433 | 7.57 | -36 | -86 | 22 |
| | | | 5.33 | -38 | -82 | 30 |
| **Conjunction analysis: scene selectivity in the SC group AND in the CC group** | | | | | | |
| R Fusiform Gyrus | 37 | 1183 | 9.15 | 24 | -32 | -18 |
| R ParaHippocampal Gyrus | 36 | | 7.81 | 32 | -42 | -8 |
| R Calcarine Gyrus | 17 | | 7.78 | 10 | -84 | 0 |
| L Calcarine Gyrus | 17 | 1082 | 9.31 | -14 | -58 | 12 |
| L Lingual Gyrus | 18 | | 7.62 | -22 | -44 | -10 |
| L ParaHippocampal Gyrus | 37 | | 7.33 | -24 | -36 | -14 |
| R Lingual Gyrus | 18 | 616 | 9.90 | 18 | -54 | 12 |
| R Calcarine Gyrus | 17 | | 8.80 | 10 | -54 | 14 |
| R Precuneus | 23 | | 4.52 | 24 | -64 | 18 |
| R Middle Occipital Gyrus | 19 | 257 | 5.74 | 44 | -76 | 22 |
| | | | 5.41 | 38 | -80 | 16 |
| | | | 4.88 | 38 | -70 | 20 |
| L Middle Occipital Gyrus | 19 | 142 | 5.49 | -34 | -82 | 22 |
| **Group comparison: scenes > other categories x SC > CC group** | | | | | | |
| L Middle Occipital Gyrus | 19 | 255 | 4.83 | -12 | -98 | -2 |

| | | | | | | |
|---|---|---|---|---|---|---|
| L Calcarine Gyrus | 17 | | 4.46 | -20 | -100 | -10 |
| L Inferior Occipital Gyrus | 19 | | 3.78 | -26 | -94 | -12 |
| L Inferior Occipital Gyrus | 19 | 119 | 4.60 | 22 | -100 | -10 |
| R Lingual Gyrus | 18 | | 4.11 | 22 | -100 | 2 |
| R Calcarine Gyrus | 17 | | 3.72 | 16 | -90 | -14 |
| **Faces vs. other categories in the SC group** | | | | | | |
| R Superior Temporal Gyrus | 22 | 5047 | 14.16 | 46 | -32 | 2 |
| R Fusiform Gyrus | 37 | | 11.97 | 52 | -22 | 0 |
| R Inferior Occipital Gyrus | 19 | | 9.01 | 38 | -80 | -14 |
| R IFG p. Triangularis | 45 | 1298 | 6.25 | 56 | 28 | 4 |
| R IFG p. Opercularis | 44 | | 6.18 | 56 | 18 | 10 |
| R Precentral Gyrus | 6 | | 5.34 | 50 | 2 | 48 |
| L Middle Temporal Gyrus | 21 | 1158 | 7.50 | -56 | -34 | 4 |
| | | | 7.36 | -56 | -50 | 8 |
| | 22 | | 6.10 | -56 | -26 | 0 |
| L Precuneus | 7 | 368 | 6.65 | 2 | -66 | 38 |
| R Precuneus | 31 | | 4.85 | 4 | -58 | 32 |
| | | | 4.61 | 4 | -54 | 24 |
| L Inferior Occipital Gyrus | 19 | 229 | 6.52 | -44 | -68 | -14 |
| L Fusiform Gyrus | 37 | | 4.86 | -40 | -86 | -12 |
| L Middle Occipital Gyrus | 19 | | 3.36 | -40 | -80 | -2 |
| L Hippocampus | | 143 | 5.08 | -34 | -2 | -14 |
| | | | 4.68 | -26 | -8 | -16 |
| R Superior Medial Gyrus | 9 | 117 | 5.33 | 8 | 62 | 28 |
| | | | 4.56 | 10 | 54 | 42 |
| L Temporal Pole | 38 | | 4.01 | -30 | 8 | -28 |
| R IFG p. Orbitalis | 47 | 82 | 4.69 | 38 | 30 | -10 |
| **Faces vs. other categories in the CC group** | | | | | | |
| R Superior Temporal Gyrus | 22 | 1650 | 6.79 | 66 | -36 | 16 |
| R Middle Temporal Gyrus | 21 | | 6.62 | 48 | -34 | -2 |
| | 22 | | 6.61 | 52 | -12 | -12 |
| R Medial Temporal Pole | 38 | 225 | 5.40 | 40 | 4 | -20 |
| | | | 4.94 | 54 | 8 | -24 |
| R Amygdala | | | 4.79 | 32 | 0 | -22 |
| R IFG p. Triangularis | 44 | 367 | 5.39 | 44 | 20 | 20 |
| | | | 5.15 | 56 | 20 | 18 |
| R IFG p. Opercularis | 44 | | 4.37 | 52 | 16 | 10 |
| R Middle Frontal Gyrus | 6 | 183 | 5.15 | 48 | 0 | 52 |
| R Precentral Gyrus | 6 | | 5.08 | 42 | 4 | 44 |
| R Middle Frontal Gyrus | 8 | | 3.79 | 34 | 2 | 36 |
| L Middle Temporal Gyrus | 21 | 80 | 4.56 | -52 | -22 | -6 |
| | | | 4.19 | -50 | -46 | 4 |
| **Conjunction analysis: face selectivity in the SC group AND in the CC group** | | | | | | |
| R Middle Temporal Gyrus | 21 | 1032 | 6.61 | 48 | -34 | 0 |
| R Superior Temporal Gyrus | 22 | | 6.48 | 54 | -36 | 8 |
| | | | 6.20 | 50 | -14 | -10 |
| R IFG p. Opercularis | 44 | 264 | 5.02 | 56 | 18 | 18 |

| | | | | | | |
|---|---|---|---|---|---|---|
| | | | 4.75 | 44 | 20 | 20 |
| R IFG p. Triangularis | 45 | | 4.25 | 54 | 22 | 2 |
| R Medial Temporal Pole | 38 | 134 | 5.40 | 40 | 4 | -20 |
| | | | 4.51 | 54 | 10 | -22 |
| R Middle Temporal Gyrus | | | 4.02 | 48 | 4 | -26 |
| R Precentral Gyrus | 6 | 87 | 4.34 | 48 | 2 | 50 |
| | | | 3.78 | 48 | 6 | 40 |
| **Group comparison: faces > other categories x SC > CC group** | | | | | | |
| R Superior Temporal Gyrus | 22 | 447 | 7.99 | 48 | -30 | 2 |
| | | | 7.56 | 52 | -22 | 0 |
| | | | 7.15 | 46 | -36 | 8 |
| R Inferior Temporal Gyrus | 37 | 207 | 5.51 | 44 | -54 | -16 |
| | | | 3.77 | 44 | -34 | -16 |
| R Middle Temporal Gyrus | 19 | 122 | 5.99 | 60 | -60 | 10 |
| | 39 | | 5.18 | 50 | -58 | 12 |
| R Inferior Occipital Gyrus | 19 | 82 | 5.43 | 38 | -80 | -14 |
| **Other objects vs. other categories in the SC group** | | | | | | |
| L IFG p. Orbitalis | 47 | 108 | 4.92 | -36 | 36 | -8 |
| | | | 4.36 | -24 | 36 | -10 |
| **Other objects vs. other categories in the CC group** | | | | | | |
| L Inferior Temporal Gyrus | 37 | 426 | 6.13 | -48 | -48 | -10 |
| | | | 6.08 | -44 | -58 | -8 |
| L Middle Temporal Gyrus | 21 | | 5.01 | -58 | -36 | -14 |
| L Middle Occipital Gyrus | 19 | 309 | 5.08 | -40 | -64 | 24 |
| | 39 | | 5.03 | -30 | -70 | 30 |
| L Inferior Parietal Lobule | 39 | | 4.13 | -30 | -74 | 42 |
| L Fusiform Gyrus | 37 | 217 | 6.26 | -36 | -14 | -20 |
| | 36 | | 5.19 | -36 | -26 | -20 |
| | | | 4.82 | -30 | -48 | -20 |
| **Conjunction analysis: other object selectivity in the SC group AND in the CC group** | | | | | | |
| *no significant clusters* | | | | | | |
| **Group comparison: other objects > other categories x SC > CC group** | | | | | | |
| *no significant clusters* | | | | | | |

*Note.* Clusters showing significantly higher activation for each of the four visually presented categories (*vs. all other categories)* selectivity-contrasts in the SC and the CC group. The voxel-wise threshold was set to $p < .001$ uncorrected and the resulting statistical maps were corrected for multiple comparisons using cluster-wise FWE-correction at $p < .05$. MNI coordinates and t-values were derived from the peak voxel of the cluster. MNI = Montreal Neurological Institute coordinates system. L = left. R = right. CC = congenital cataract reversal individuals. SC = normally-sighted individuals. IFG = inferior frontal gyrus.

**Table S3.** Brain regions with significant clusters in category-selective contrasts in the SC and the CC group for the auditory runs.

| Contrast | Hemisphere | Brain Region | Brodmann Area | Cluster size | t Statistics | MNI coordinates x | y | z |
|---|---|---|---|---|---|---|---|---|
| **Bodies vs. other categories in the SC group** | | | | | | | | |
| | *no significant clusters* | | | | | | | |
| **Bodies vs. other categories in the CC group** | | | | | | | | |
| | L Posterior-Medial Frontal | | 6 | 328 | 6.23 | -2 | 2 | 68 |
| | | | | | 4.72 | -8 | 4 | 58 |
| | | | | | 4.20 | 12 | 0 | 72 |
| | R IFG p. Opercularis | | 44 | 264 | 5.40 | 60 | 14 | 20 |
| | R Precentral Gyrus | | 6 | | 5.22 | 46 | 4 | 38 |
| | | | | | 4.32 | 44 | 4 | 50 |
| | L Middle Temporal Gyrus | | 21 | 165 | 6.35 | -48 | -50 | 8 |
| | | | | | 4.53 | -40 | -44 | 8 |
| | L Middle Temporal Gyrus | | 37 | | 4.44 | -54 | -54 | -4 |
| | L IFG p. Opercularis | | 44 | 125 | 4.39 | -44 | 6 | 20 |
| | | | | | 4.38 | -58 | 14 | 26 |
| | L Precentral Gyrus | | 6 | | 4.05 | -60 | 6 | 32 |
| | R Superior Temporal Gyrus | | 22 | 112 | 4.19 | 46 | -40 | 8 |
| | | | | | 3.96 | 60 | -32 | 10 |
| | | | | | 3.93 | 54 | -38 | 14 |
| | L Superior Temporal Gyrus | | 22 | 80 | 4.82 | -54 | -38 | 14 |
| | | | | | 3.74 | -64 | -44 | 14 |
| **Conjunction analysis: body selectivity in the SC group AND in the CC group** | | | | | | | | |
| | *no significant clusters* | | | | | | | |
| **Group comparison: bodies > other categories x SC > CC group** | | | | | | | | |
| | *no significant clusters* | | | | | | | |
| **Scenes vs. other categories in the SC group** | | | | | | | | |
| | *no significant clusters* | | | | | | | |
| **Scenes vs. other categories in the CC group** | | | | | | | | |
| | L Fusiform Gyrus | | 37 | 107 | 5.65 | -34 | -32 | -16 |
| | | | | | 4.47 | -28 | -38 | -18 |
| | L Precuneus | | 23 | 90 | 4.84 | -10 | -56 | 12 |
| | L Calcarine Gyrus | | 17 | | 4.50 | -14 | -52 | 4 |
| | | | | | 3.62 | -4 | -50 | 8 |
| **Conjunction analysis: scene selectivity in the SC group AND in the CC group** | | | | | | | | |
| | *no significant clusters* | | | | | | | |
| **Group comparison: scenes > other categories x SC > CC group** | | | | | | | | |
| | *no significant clusters* | | | | | | | |
| **Faces vs. other categories in the SC group** | | | | | | | | |
| | R Superior Temporal Gyrus | | 22 | 2249 | 10.1 | 60 | -2 | -2 |

| | | | | | | |
|---|---|---|---|---|---|---|
| | | | 9.95 | 58 | -32 | 2 |
| | | | 8.82 | 60 | -24 | 0 |
| L Middle Temporal Gyrus | 22 | 1207 | 8.07 | -66 | -26 | 4 |
| | | | 7.65 | -62 | -20 | 0 |
| L Superior Temporal Gyrus | 22 | | 7.55 | -58 | -10 | 0 |
| L Temporal Pole | 38 | 261 | 5.76 | -52 | 12 | -14 |
| | | | 5.23 | -50 | 14 | -22 |
| | 22 | | 4.80 | -56 | 10 | -6 |
| L Angular Gyrus | 39 | 208 | 4.25 | -40 | -70 | 48 |
| | | | 4.18 | -40 | -68 | 38 |
| L Inferior Parietal Lobule | 39 | | 3.64 | -34 | -60 | 42 |
| L Precuneus | 7 | 396 | 4.54 | -2 | -52 | 52 |
| | 31 | | 4.51 | -10 | -58 | 42 |
| | 7 | | 4.20 | -6 | -62 | 52 |
| L Middle Frontal Gyrus | 8 | 81 | 5.19 | -26 | 20 | 46 |
| **Faces vs. other categories in the CC group** | | | | | | |
| R Middle Temporal Gyrus | 21 | 1144 | 7.49 | 50 | -20 | -12 |
| R Superior Temporal Gyrus | 22 | | 7.23 | 60 | -14 | -8 |
| | | | 6.42 | 62 | -2 | -2 |
| L Middle Temporal Gyrus | 22 | 1089 | 7.61 | -62 | -8 | -6 |
| | | | 6.06 | -64 | -18 | -4 |
| | | | 5.89 | -54 | -18 | -6 |
| L Superior Medial Gyrus | 9 | 141 | 4.99 | 0 | 62 | 26 |
| R Superior Medial Gyrus | 9 | | 3.86 | 8 | 56 | 24 |
| R Medial Temporal Pole | 38 | 107 | 5.47 | 52 | 6 | -24 |
| R Temporal Pole | | | 4.14 | 40 | 4 | -22 |
| **Conjunction analysis: face selectivity in the SC group AND in the CC group** | | | | | | |
| R Superior Temporal Gyrus | 22 | 800 | 6.68 | 62 | -18 | -4 |
| | | | 6.42 | 62 | -2 | -2 |
| R Temporal Pole | 22 | | 5.74 | 58 | 6 | -10 |
| L Superior Temporal Gyrus | 22 | 486 | 6.79 | -60 | -6 | -2 |
| L Middle Temporal Gyrus | 22 | | 5.46 | -58 | -16 | -2 |
| | 21 | | 5.23 | -66 | -24 | -4 |
| L Temporal Pole | 38 | 98 | 5.11 | -52 | 12 | -18 |
| | | | 4.47 | -56 | 8 | -12 |
| | | | 3.59 | -46 | 2 | -18 |
| **Group comparison: faces > other categories x SC > CC group** | | | | | | |
| R Superior Temporal Gyrus | 22 | 116 | 5.57 | 60 | -32 | 4 |
| | | | 4.03 | 50 | -28 | 2 |
| | | | 3.69 | 60 | -24 | 0 |
| **Other objects vs. other categories in the SC group** | | | | | | |
| L Inferior Temporal Gyrus | 37 | 116 | 4.62 | -50 | -48 | -18 |
| | | | 4.59 | -44 | -32 | -14 |
| | | | 4.53 | -40 | -40 | -14 |
| L IFG p. Orbitalis | 47 | 115 | 7.16 | -34 | 36 | -8 |
| | | | 4.22 | -24 | 36 | -10 |
| L Middle Temporal Gyrus | 37 | 108 | 5.01 | -42 | -58 | -4 |
| L Inferior Occipital Gyrus | 19 | | 4.20 | -46 | -66 | -4 |

**Other objects vs. other categories in the CC group**

*no significant clusters*

**Conjunction analysis: body selectivity in the SC group AND in the CC group**

*no significant clusters*

**Group comparison: other objects > other categories x SC > CC group**

*no significant clusters*

*Note.* Clusters showing significantly higher activation for each of the four auditorily presented categories (*vs. all other categories)* selectivity-contrasts in the SC and the CC group The voxel-wise threshold was set to $p < .001$ uncorrected and the resulting statistical maps were corrected for multiple comparisons using cluster-wise FWE-correction at $p < .05$. MNI coordinates and t-values are derived from the peak voxel of the cluster. MNI = Montreal Neurological Institute coordinates system. L = left. R = right. CC = congenital cataract reversal individuals. SC = normally-sighted individuals. IFG = inferior frontal gyrus.

**Table S4.** Whole-brain searchlight analysis for visual and auditory categories in the SC and the CC group.

| Contrast | Hemisphere | Brain Region | Brodmann Area | Cluster size | t Statistics | MNI coordinates x | y | z |
|---|---|---|---|---|---|---|---|---|
| **searchlight decoding for visual categories (*bodies, scenes, faces* and *other objects*)** | | | | | | | | |
| **the SC group** | | | | | | | | |
| | L Calcarine Gyrus | | 17 | 17566 | 12.79 | -6 | -90 | -2 |
| | | | | | 12.46 | -14 | -96 | -2 |
| | R Fusiform Gyrus | | 37 | | 12.29 | 34 | -42 | -18 |
| | L IFG p. Opercularis | | 44 | 350 | 8.87 | -38 | 8 | 26 |
| | L Precentral Gyrus | | 8 | | 8.17 | -44 | 12 | 30 |
| | | | 6 | | 8.08 | -44 | 4 | 34 |
| | R Middle Frontal Gyrus | | 6 | 291 | 10.33 | 38 | 0 | 56 |
| | R Precentral Gyrus | | 6 | | 10.21 | 48 | -10 | 48 |
| | | | 4 | | 8.24 | 40 | -16 | 54 |
| | R Superior Temporal Gyrus | | 22 | 280 | 7.90 | 52 | 2 | -14 |
| | | | | | 7.22 | 54 | 0 | -6 |
| | R Temporal Pole | | | | 7.17 | 58 | 18 | -8 |
| | R Medial Temporal Pole | | 38 | 170 | 9.40 | 42 | 6 | -38 |
| | | | 20 | | 7.23 | 42 | 2 | -48 |
| | R Inferior Temporal Gyrus | | 20 | | 6.30 | 50 | 0 | -38 |
| | R Middle Frontal Gyrus | | 9 | 192 | 7.99 | 42 | 32 | 22 |
| | R IFG p. Opercularis | | 44 | | 6.43 | 40 | 14 | 32 |
| | R Middle Frontal Gyrus | | 9 | | 6.30 | 38 | 24 | 32 |
| | L SupraMarginal Gyrus | | 40 | 147 | 7.40 | -48 | -44 | 24 |
| | | | | | 6.80 | -42 | -44 | 30 |
| | | | | | 6.07 | -50 | -32 | 30 |
| | R Cuneus | | 18 | 137 | 6.59 | 20 | -58 | 20 |
| | R Calcarine Gyrus | | 17 | | 6.13 | 16 | -56 | 6 |
| | | | | | 6.06 | 16 | -62 | 14 |
| | L Middle Temporal Gyrus | | 21 | 132 | 8.67 | -56 | -30 | -8 |
| | | | | | 6.54 | -58 | -32 | 0 |
| | | | | | 5.20 | -50 | -42 | 6 |
| | R Posterior-Medial Frontal | | 8 | 118 | 8.20 | 4 | 16 | 46 |
| | R MCC | | 8 | | 8.14 | 10 | 20 | 40 |
| | L Posterior-Medial Frontal | | 8 | | 6.49 | -2 | 22 | 46 |
| | L Precuneus | | 31 | 108 | 7.51 | -10 | -44 | 46 |
| | R Precuneus | | 31 | | 6.31 | 4 | -42 | 48 |
| | L Precuneus | | 23 | 72 | 6.17 | -12 | -58 | 14 |
| | L Calcarine Gyrus | | 17 | | 5.73 | -16 | -60 | 6 |
| | L SupraMarginal Gyrus | | 39 | 71 | 7.69 | -58 | -48 | 30 |
| | L SupraMarginal Gyrus | | 39 | | 5.17 | -60 | -54 | 24 |
| | R Posterior-Medial Frontal | | 6 | 70 | 6.61 | 4 | -10 | 78 |
| | | | | | 6.29 | 2 | -4 | 64 |
| | R Posterior-Medial Frontal | | 6 | 51 | 8.01 | 6 | 6 | 58 |
| | | | | | 5.34 | 2 | 0 | 52 |
| | R Precentral Gyrus | | 6 | 36 | 5.96 | 50 | 4 | 34 |
| | L Superior Temporal Gyrus | | 22 | 36 | 6.13 | -52 | -4 | -4 |
| | R Precuneus | | 31 | 33 | 9.52 | 14 | -48 | 48 |
| | R Precuneus | | 31 | 29 | 6.11 | 4 | -64 | 24 |
| | L Postcentral Gyrus | | 1 | 28 | 6.43 | -56 | -20 | 26 |
| | L IFG p. Orbitalis | | 47 | 22 | 5.97 | -40 | 36 | -4 |

| | | | | | | | |
|---|---|---|---|---|---|---|---|
| **the CC group** | | | | | | | |
| | R Middle Temporal Gyrus | 37 | 8878 | 11.42 | 54 | -60 | 2 |
| | L Lingual Gyrus | 18 | | 11.31 | -18 | -58 | -12 |
| | R Calcarine Gyrus | 17 | | 10.34 | 16 | -78 | 16 |
| | R Middle Frontal Gyrus | 9 | 391 | 8.24 | 40 | 36 | 18 |
| | R IFG p. Triangularis | 9 | | 7.96 | 50 | 24 | 20 |
| | | 44 | | 7.16 | 54 | 16 | 22 |
| | L IFG p. Opercularis | 44 | 354 | 7.65 | -38 | 8 | 26 |
| | L Middle Frontal Gyrus | 8 | | 7.29 | -48 | 22 | 36 |
| | L Precentral Gyrus | | | 6.80 | -54 | 12 | 30 |
| | R SupraMarginal Gyrus | 40 | 255 | 6.98 | 62 | -28 | 30 |
| | | | | 6.51 | 54 | -20 | 24 |
| | L Precentral Gyrus | 6 | 189 | 9.53 | -40 | 4 | 44 |
| | | | | 7.90 | -46 | 0 | 48 |
| | R Fusiform Gyrus | 37 | 168 | 7.95 | 34 | -42 | -18 |
| | | | | 7.18 | 38 | -50 | -16 |
| | R Hippocampus | | | 6.70 | 28 | -32 | -8 |
| | L Superior Frontal Gyrus | 6 | 131 | 7.75 | -18 | 18 | 60 |
| | L Posterior-Medial Frontal | 6 | | 7.15 | -10 | 22 | 60 |
| | L Middle Frontal Gyrus | 6 | | 6.74 | -28 | 12 | 52 |
| | L Superior Medial Gyrus | 8 | 97 | 7.60 | 0 | 28 | 54 |
| | L Posterior-Medial Frontal | 8 | | 6.48 | -2 | 22 | 46 |
| | R Middle Frontal Gyrus | 6 | 74 | 8.56 | 38 | -2 | 56 |
| | Hippocampus | 36 | 68 | 8.64 | -32 | -22 | -22 |
| | R Lingual Gyrus | 19 | 60 | 5.93 | 22 | -54 | -8 |
| | R Cerebelum | | | 5.72 | 28 | -50 | -20 |
| | L Inferior Temporal Gyrus | 37 | 57 | 6.26 | -42 | -50 | -16 |
| | | | | 5.80 | -52 | -50 | -18 |
| | R Calcarine Gyrus | 17 | 53 | 6.40 | 20 | -56 | 18 |
| | | | | 5.29 | 16 | -58 | 10 |
| | L Middle Temporal Gyrus | 21 | 47 | 7.09 | -64 | -8 | -20 |
| | | | | 5.53 | -56 | -10 | -22 |
| | R Superior Temporal Gyrus | 41 | | 6.25 | 60 | -24 | 12 |
| | L Posterior-Medial Frontal | 6 | 45 | 7.39 | -4 | -2 | 70 |
| | R Fusiform Gyrus | 19 | 38 | 6.89 | 30 | -76 | -12 |
| | L Calcarine Gyrus | 23 | 36 | 6.00 | -8 | -60 | 10 |
| | R Cerebelum | | 30 | 6.26 | 6 | -82 | -26 |
| | | | 27 | 7.83 | -6 | -66 | -6 |
| | R Precuneus | 23 | 24 | 6.21 | 8 | -50 | 12 |
| | L Precuneus | 7 | 23 | 5.99 | -2 | -46 | 60 |
| | R Precuneus | 31 | 22 | 6.18 | 4 | -66 | 22 |
| **Group comparison: SC > CC** | | | | | | | |
| | L Calcarine Gyrus | 17 | 70 | 6.25 | -6 | -90 | -2 |
| | | | | 6.08 | -14 | -96 | -2 |
| **searchlight decoding for auditory categories (*bodies, scenes, faces* and *other objects*)** | | | | | | | |
| **the SC group** | | | | | | | |
| | R Superior Temporal Gyrus | 22 | 2151 | 13.67 | 58 | -18 | 0 |
| | | 41 | | 13.32 | 66 | -18 | 8 |
| | | 22 | | 9.79 | 58 | -32 | 2 |
| | L Superior Temporal Gyrus | 22 | 601 | 9.14 | -56 | -2 | -4 |

| | | | | | | |
|---|---|---|---|---|---|---|
| | | | 9.09 | -48 | -16 | -2 |
| | | | 8.71 | -64 | -26 | 4 |
| R Precuneus | 7 | 143 | 12.01 | 6 | -60 | 44 |
| L Precuneus | 31 | | 6.70 | -4 | -60 | 42 |
| R Precuneus | 7 | | 6.56 | 2 | -66 | 40 |
| L Superior Temporal Gyrus | 22 | 130 | 9.99 | -50 | -30 | 4 |
| L Middle Temporal Gyrus | 22 | | 6.86 | -52 | -40 | 8 |
| L Superior Temporal Gyrus | 22 | | 5.80 | -52 | -44 | 16 |
| L Middle Temporal Gyrus | 39 | 90 | 7.69 | -54 | -58 | 12 |
| | | | 6.94 | -54 | -64 | 20 |
| R Angular Gyrus | 39 | 54 | 7.60 | 40 | -62 | 34 |
| | | | 5.77 | 38 | -64 | 44 |
| | | 54 | 6.41 | 42 | -42 | 34 |
| R Inferior Parietal Lobule | 7 | | 5.93 | 38 | -42 | 46 |
| R Middle Occipital Gyrus | 19 | 37 | 6.80 | 32 | -68 | 34 |
| | | | 5.87 | 32 | -74 | 40 |
| L Middle Frontal Gyrus | | 29 | 7.45 | -32 | 14 | 46 |
| L Angular Gyrus | 39 | 22 | 6.90 | -40 | -64 | 36 |
| | 39 | | 5.90 | 34 | -48 | 40 |
| R Cerebelum | | 20 | 8.09 | 12 | -76 | -32 |
| **the CC group** | | | | | | |
| L Superior Temporal Gyrus | 40 | 1407 | 12.69 | -54 | -26 | 12 |
| | | | 11.64 | -46 | -32 | 20 |
| L Middle Temporal Gyrus | 22 | | 9.18 | -64 | -28 | 4 |
| R Superior Temporal Gyrus | 22 | 1111 | 9.24 | 58 | -18 | 0 |
| | 41 | | 8.37 | 66 | -22 | 8 |
| R Rolandic Operculum | 40 | | 8.05 | 52 | -16 | 12 |
| L Superior Temporal Gyrus | 22 | 198 | 7.60 | -54 | -2 | -4 |
| L Temporal Pole | 22 | | 6.85 | -52 | 8 | -4 |
| L Rolandic Operculum | 6 | | 6.62 | -52 | 2 | 6 |
| R IFG p. Triangularis | 45 | 136 | 7.03 | 58 | 26 | 28 |
| | | | 6.87 | 56 | 24 | 6 |
| | 46 | | 6.52 | 46 | 34 | 12 |
| L Superior Parietal Lobule | | 61 | 6.69 | -14 | -76 | 56 |
| L Precuneus | 7 | | 6.46 | -8 | -78 | 50 |
| L Superior Parietal Lobule | 7 | | 5.07 | -20 | -78 | 48 |
| L Precentral Gyrus | 6 | 49 | 8.29 | -48 | 10 | 44 |
| L Superior Temporal Gyrus | 22 | 45 | 10.27 | -48 | -16 | -2 |
| | | | 6.83 | -40 | -8 | -10 |
| L Angular Gyrus | 39 | 31 | 6.79 | -48 | -70 | 36 |
| L Angular Gyrus | 39 | 24 | 5.65 | -40 | -66 | 40 |
| L IFG p. Triangularis | 44 | 23 | 8.28 | -52 | 20 | 16 |
| L IFG p. Triangularis | 46 | | 5.38 | -52 | 28 | 18 |
| R Middle Occipital Gyrus | 19 | 23 | 7.00 | 32 | -68 | 38 |
| R IFG p. Opercularis | 44 | 22 | 6.37 | 58 | 10 | 26 |
| R Precentral Gyrus | 6 | | 6.29 | 50 | 8 | 34 |
| **Group comparison: SC > CC** | | | | | | |
| *no significant clusters* | | | | | | |

*Note.* Clusters showing high classifier performance in whole brain searchlight analysis for the visual and auditory categories in the SC and the CC group. The voxel-wise threshold was set to $p < .0001$ uncorrected and the resulting statistical maps were corrected for multiple comparisons using cluster-wise FWE-correction at $p < .05$. MNI coordinates and t-values are derived from the peak voxel of the cluster. MNI = Montreal Neurological Institute coordinates system. L = left. R = right. CC = congenital cataract reversal individuals. SC = normally-sighted individuals. IFG = inferior frontal gyrus.

**Table S5**. Within-subject classification results for visual and auditory categories in the SC and the CC group.

| **ROI name** | | **SC group** mean | **SC group** SEM | **SC group** vs chance t statistics | **CC group** mean | **CC group** SEM | **CC group** vs chance t statistics | **SC vs CC group** t statistics |
|---|---|---|---|---|---|---|---|---|
| **early visual cortex** | visual categories | 70.31 | 5.03 | 8.42*** | 60.94 | 6.32 | 5.32*** | 1.09 |
| | auditory categories | 28.91 | 2.46 | 3.42 | 28.13 | 3.49 | 4.02 | 0.48 |
| **VOTC** | visual categories | 91.41 | 3.47 | 25.30*** | 75.00 | 6.99 | 6.69*** | 2.07$^{t}$ |
| | auditory categories | 32.03 | 3.39 | 4.41 | 35.16 | 2.19 | 2.61 | -0.36 |
| **primary auditory cortex** | visual categories | 44.53 | 2.34 | 6.52*** | 40.63 | 3.31 | 4.41** | 0.84 |
| | auditory categories | 36.72 | 3.21 | 1.49 | 39.06 | 3.27 | 0.84 | -0.17 |
| **higher auditory cortex** | visual categories | 43.75 | 3.31 | 5.29*** | 45.31 | 4.38 | 4.33*** | -0.27 |
| | auditory categories | 41.41 | 3.48 | 1.94 | 39.06 | 5.03 | 4.33 | 0.72 |

*Note.* Classification accuracy was averaged over four categories (*bodies, scenes, faces* and *other objects*) for visual and auditory categories and is shown in %. Chance level = 25%. Thresholds: *p < .05, **p < .01, ***p < .001, *t* = trend level of p = .057. SC = normally-sighted individuals. CC = congenital cataract reversal individuals. SEM = standard error of the mean. VOTC = ventral occipito-temporal cortex.

**Table S6**. Classification accuracy for single categories in within-subject classification for visual and auditory categories in the SC and the CC group.

| ROI name | | | visual categories body | scene | face | object | auditory categories body | scene | face | object |
|---|---|---|---|---|---|---|---|---|---|---|
| early visual cortex | SC group | mean | 81.25 | 62.50 | 65.63 | 71.88 | 21.88 | 28.13 | 25.00 | 31.25 |
| | | SEM | 8.56 | 6.25 | 6.15 | 8.19 | 6.90 | 5.30 | 7.65 | 7.33 |
| | vs. chance | t statistics | 6.89*** | 6.30*** | 6.92*** | 6.02*** | -1.02 | 1.12 | 0.71 | 1.33 |
| | CC group | mean | 65.63 | 56.25 | 75.00 | 43.75 | 18.75 | 34.38 | 37.50 | 18.75 |
| | | SEM | 9.82 | 10.60 | 8.84 | 7.33 | 5.85 | 8.77 | 8.84 | 5.85 |
| | vs. chance | t statistics | 4.42** | 3.25* | 5.95*** | 2.87* | -1.51 | 1.51 | 1.81 | -1.51 |
| | SC vs. CC | t statistics | 1.56 | 1.03 | -1.30 | 2.75* | 0.92 | -1.10 | -1.46 | 1.67 |
| VOTC | SC group | mean | 87.50 | 93.75 | 96.88 | 87.50 | 25.00 | 18.75 | 40.63 | 40.63 |
| | | SEM | 4.42 | 3.83 | 2.92 | 6.25 | 7.65 | 7.33 | 7.57 | 8.77 |
| | vs. chance | t statistics | 14.652*** | 18.59*** | 25.42*** | 10.39*** | 0.71 | -1.33 | 2.40* | 2.14 |
| | CC group | mean | 75.00 | 87.50 | 68.75 | 68.75 | 9.38 | 34.38 | 43.75 | 50.00 |
| | | SEM | 8.84 | 6.25 | 8.56 | 9.63 | 4.28 | 6.15 | 8.56 | 10.83 |
| | vs. chance | t statistics | 5.95*** | 10.39*** | 5.40*** | 4.83** | -3.94 | 1.90 | 2.52* | 2.63* |
| | SC vs. CC | t statistics | 1.62 | 1.29 | 3.26** | 1.93 | 2.06 | -1.93 | -0.87 | -1.15 |
| primary auditory cortex | SC group | mean | 46.88 | 46.88 | 43.75 | 37.50 | 31.25 | 28.13 | 46.88 | 37.50 |
| | | SEM | 6.90 | 8.19 | 5.85 | 7.65 | 7.33 | 9.31 | 8.19 | 7.65 |
| | vs. chance | t statistics | 3.46** | 2.98* | 3.50** | 2.00 | 1.33 | 0.93 | 2.98* | 2.00 |
| | CC group | mean | 46.88 | 31.25 | 34.38 | 50.00 | 15.63 | 43.75 | 46.88 | 50.00 |
| | | SEM | 6.90 | 9.63 | 6.15 | 6.25 | 8.77 | 7.33 | 8.19 | 6.25 |
| | vs. chance | t statistics | 3.47** | 1.17 | 1.90 | 4.29** | -1.51 | 2.87* | 2.98* | 4.29** |
| | SC vs. CC | t statistics | 0.69 | 1.59 | 1.49 | -1.62 | 1.70 | -1.66 | 0.69 | -1.62 |
| higher auditory cortex | SC group | mean | 37.50 | 46.88 | 50.00 | 37.50 | 53.13 | 34.38 | 50.00 | 28.13 |
| | | SEM | 4.42 | 6.90 | 7.65 | 7.65 | 5.30 | 7.57 | 9.88 | 8.19 |
| | vs. chance | t statistics | 3.13* | 3.46** | 3.56** | 2.00 | 5.60*** | 1.65 | 2.84* | 0.97 |
| | CC group | mean | 34.38 | 43.75 | 50.00 | 46.88 | 34.38 | 40.63 | 53.13 | 25.00 |
| | | SEM | 8.77 | 8.56 | 4.42 | 8.19 | 8.77 | 8.77 | 6.90 | 8.84 |
| | vs. chance | t statistics | 1.51 | 2.52* | 5.96*** | 2.98* | 1.51 | 2.14 | 4.36** | 0.71 |
| | SC vs. CC | t statistics | 0.90 | 0.87 | 0.69 | -1.27 | 2.10 | -1.05 | -0.86 | 0.86 |

*Note.* Mean classification accuracy for each category *(vs. all other categories)* was derived from the unbiased confusion matrix in the within-subject classification and is shown in %. Each value was tested against chance level (= 25%) with one sample t-test and between the groups in each ROI with two-sample t-tests. Thresholds: *p < .05, **p < .01, ***p < .001. SC = normally-sighted individuals. CC = congenital cataract reversal individuals. SEM = standard error of the mean. VOTC = ventral occipito-temporal cortex.

**Table S7**. Cross-modal classification results in the SC and the CC group.

| ROI name | decoding | SC group mean | SC group SEM | SC group vs chance t statistics | CC group mean | CC group SEM | CC group vs chance t statistics | SC vs CC group t statistics |
|---|---|---|---|---|---|---|---|---|
| **early visual cortex** | visual to auditory | 24.22 | 2.05 | -0.36 | 25.78 | 0.73 | 1.00 | -0.67 |
| | auditory to visual | 24.22 | 2.33 | -0.31 | 24.22 | 0.73 | -1.00 | 0.00 |
| **VOTC** | visual to auditory | 28.91 | 1.54 | 2.38* | 26.56 | 1.46 | 1.00 | 1.03 |
| | auditory to visual | 27.34 | 1.54 | 1.43 | 28.91 | 1.89 | 1.93* | -0.60 |
| **primary auditory cortex** | visual to auditory | 30.47 | 2.80 | 1.82 | 32.03 | 2.33 | 2.83* | -0.40 |
| | auditory to visual | 27.34 | 3.30 | 0.66 | 31.25 | 3.31 | 1.76 | -0.78 |
| **higher auditory cortex** | visual to auditory | 31.25 | 2.92 | 2.00* | 35.16 | 2.91 | 3.26** | -0.89 |
| | auditory to visual | 22.66 | 1.89 | -1.16 | 32.03 | 1.32 | 4.97*** | -3.79** |

*Note.* Classification accuracy was averaged over four categories (*bodies, scenes, faces* and *other objects*) in visual-to-auditory and auditory-to-visual decoding and is shown in %. Chance level = 25%. Thresholds: *p < .05, **p < .01, ***p < .001. SC = normally-sighted individuals. CC = congenital cataract reversal individuals. SEM = standard error of the mean. VOTC = ventral occipito-temporal cortex.

**Table S8**. Classification accuracy for single categories in cross-modal decoding in the SC and the CC group.

| ROI name | | | visual to auditory decoding body | scene | face | object | auditory to visual decoding body | scene | face | object |
|---|---|---|---|---|---|---|---|---|---|---|
| **early visual cortex** | SC group | mean | 37.50 | 6.25 | 40.63 | 12.50 | 3.13 | 34.38 | 28.13 | 28.13 |
| | | SEM | 15.31 | 3.83 | 15.27 | 6.25 | 2.92 | 13.21 | 13.58 | 14.95 |
| | vs. chance | t statistics | 1.30 | -5.19** | 1.47 | -2.34 | -7.81*** | 1.21 | 0.86 | 0.85 |
| | CC group | mean | 0.00 | 15.63 | 68.75 | 18.75 | 18.75 | 12.50 | 21.88 | 43.75 |
| | | SEM | 0.00 | 0.24 | 0.01 | 0.31 | 9.63 | 11.69 | 13.58 | 15.78 |
| | vs. chance | t statistics | - | -1.29 | 3.32* | -1.09 | -1.17 | -1.51 | -0.86 | 1.61 |
| | SC vs. CC | t statistics | 2.65* | 1.22 | 1.68 | 1.01 | 1.86 | 1.60 | 0.90 | 1.18 |
| **VOTC** | SC group | mean | 0.00 | 62.50 | 31.25 | 21.88 | 12.50 | 21.88 | 25.00 | 50.00 |
| | | SEM | 0.00 | 14.66 | 14.49 | 11.21 | 11.69 | 12.05 | 15.31 | 15.93 |
| | vs. chance | t statistics | - | 2.87* | 1.00 | -0.89 | -1.51 | -0.88 | 0.71 | 1.95 |
| | CC group | mean | 9.38 | 37.50 | 12.50 | 43.75 | 31.25 | 37.50 | 15.63 | 31.25 |
| | | SEM | 6.15 | 17.12 | 11.69 | 16.39 | 14.49 | 14.66 | 6.15 | 14.49 |
| | vs. chance | t statistics | -2.85 | 1.23* | -1.51 | 1.57 | 1.00 | 1.33 | -1.90 | 1.00 |
| | SC vs. CC | t statistics | -1.83 | 1.49 | 1.41 | -1.48 | -1.41 | -1.26 | 1.07 | 1.30 |
| **primary auditory cortex** | SC group | mean | 15.63 | 56.25 | 9.38 | 40.63 | 21.88 | 31.25 | 15.63 | 40.63 |
| | | SEM | 11.64 | 15.78 | 8.77 | 10.77 | 10.31 | 13.07 | 7.57 | 11.64 |
| | vs. chance | t statistics | -1.29 | 2.32t | -2.14 | 1.84 | -0.91 | 1.04 | -1.65 | 1.74 |
| | CC group | mean | 21.88 | 56.25 | 18.75 | 25.00 | 53.13 | 21.88 | 34.38 | 15.63 |
| | | SEM | 9.31 | 12.30 | 9.63 | 13.26 | 10.31 | 8.19 | 12.45 | 7.57 |
| | vs. chance | t statistics | -0.93 | 2.85* | -1.17 | 0.71 | 3.04 | -0.97 | 1.25 | -1.65 |
| | SC vs. CC | t statistics | -0.97 | 0.69 | -1.18 | 1.33 | -2.38 | 1.10 | -1.63 | 2.07 |
| **higher auditory cortex** | SC group | mean | 12.50 | 50.00 | 9.38 | 53.13 | 34.38 | 21.88 | 6.25 | 25.00 |
| | | SEM | 8.84 | 10.83 | 6.15 | 8.19 | 13.93 | 8.19 | 5.85 | 9.88 |
| | vs. chance | t statistics | -1.81 | 2.63* | -2.85* | 3.72** | 1.19 | -0.97 | -3.50** | 0.71 |
| | CC group | mean | 34.38 | 43.75 | 34.38 | 28.13 | 56.25 | 34.38 | 15.63 | 21.88 |
| | | SEM | 9.82 | 14.49 | 12.45 | 10.31 | 10.60 | 10.77 | -6.15 | -8.19 |
| | vs. chance | t statistics | 1.41 | 1.70 | 1.25 | 0.91 | 3.25* | 1.34 | 1.90 | 0.97 |
| | SC vs. CC | t statistics | -1.95 | 0.92 | -2.07 | 2.16* | -1.60 | -1.34 | -1.49 | 0.85 |

*Note.* Mean classification accuracy for each category *(vs. all other categories)* was derived from the unbiased confusion matrix in the cross-modal classification and is shown in %. Each value was tested against chance level (= 25%) with one sample t-test and between the groups in each ROI with two-sample t-tests. Thresholds: *p < .05, **p < .01, ***p < .001. SC = normally-sighted individuals. CC = congenital cataract reversal individuals. SEM = standard error of the mean. VOTC = ventral occipito-temporal cortex.

**Table S9.** Summary of statistical analyses (mixed-design ANOVA) for category-selectivity between normally-sighted individuals and congenital cataract reversal individuals of the main ROI analysis (Fig. 2).

| **ROI** | | H | Group | | | Category | | | | Group x Condition Interaction | | | |
|---|---|---|---|---|---|---|---|---|---|---|---|---|---|
| | | | F(1,14) | Sig. | Effect size (eta$^2$) | F(3,42) | Sig. | | Effect size (eta$^2$) | F(3,42) | Sig. | | Effect size (eta$^2$) |
| **faces** | IOG | L | 0.02 | .890 | 0.001 | 25.60 | **<.001** | * | 0.135 | 0.90 | .114 | * | 0.005 |
| | | R | 4.82 | **.045** | 0.182 | 49.63 | **<.001** | | 0.557 | 4.10 | **.012** | | 0.094 |
| | mFus | L | 0.01 | .921 | 0.000 | 12.83 | **<.001** | * | 0.239 | 1.21 | .308 | * | 0.029 |
| | | R | 1.26 | .281 | 0.059 | 45.06 | **<.001** | * | 0.489 | 13.60 | **<.001** | * | 0.224 |
| | pFus | L | 0.52 | .482 | 0.029 | 19.25 | **<.001** | * | 0.213 | 1.17 | .317 | * | 0.016 |
| | | R | 0.41 | .531 | 0.024 | 78.29 | **<.001** | | 0.497 | 14.11 | **<.001** | | 0.151 |
| **places** | COS | L | 0.68 | .425 | 0.032 | 37.93 | **<.001** | * | 0.467 | 2.45 | .077 | * | 0.054 |
| | | R | 0.32 | .580 | 0.018 | 74.81 | **<.001** | * | 0.522 | 4.11 | **.037** | * | 0.056 |
| | TOS | R | 0.10 | .754 | 0.005 | 53.59 | **<.001** | * | 0.546 | 0.49 | .593 | * | 0.011 |
| **bodies** | ITG | L | 2.26 | .155 | 0.102 | 43.85 | **<.001** | * | 0.481 | 3.83 | .057 | * | 0.075 |
| | | R | 2.17 | .162 | 0.108 | 121.30 | **<.001** | * | 0.660 | 7.45 | **.008** | * | 0.107 |
| | LOS | L | 1.94 | .185 | 0.098 | 40.77 | **<.001** | * | 0.391 | 6.06 | **.016** | * | 0.087 |
| | | R | 0.54 | .475 | 0.031 | 47.88 | **<.001** | * | 0.352 | 6.82 | **.005** | * | 0.072 |
| | MTG | L | 1.24 | .285 | 0.065 | 54.18 | **<.001** | * | 0.453 | 0.60 | .524 | * | 0.009 |
| | | R | 0.84 | .074 | 0.040 | 68.41 | **<.001** | * | 0.598 | 1.37 | .271 | * | 0.029 |
| | OTS | L | 0.04 | .838 | 0.002 | 11.19 | **.002** | * | 0.226 | 1.25 | .293 | * | 0.031 |
| | | R | 1.30 | .273 | 0.064 | 36.59 | **<.001** | | 0.411 | 7.50 | **<.001** | | 0.125 |
| **other objects** | LOC | L | 0.19 | .670 | 0.011 | 5.34 | **.011** | * | 0.052 | 0.23 | .800 | * | 0.002 |
| | | R | 1.96 | .184 | 0.093 | 13.92 | **<.001** | * | 0.207 | 1.54 | .236 | * | 0.028 |

*Note*. *Greenhouse–Geisser corrected for lack of sphericity. H = hemisphere. L = Left. R = Right. Sig = Statistical significance. ROI = region of interest, for full description of ROIs see visual atlas of Rosenke et al. (2021). LOC = Lateral Occipital Complex.

**Table S10.** Summary of statistical analyses (repeated-measures ANOVA) separately in normally-sighted individuals (the SC group) and in congenital cataract reversal individuals (the CC group) of the main ROI analysis (Fig. 2).

| ROI | | H | The SC Group | | | The CC Group | | |
|---|---|---|---|---|---|---|---|---|
| | | | F(3,21) | Sig. | Effect size (eta$^2$) | F(3,21) | Sig. | Effect size (eta$^2$) |
| faces | IOG | L | 12.32 | **<.001** | 0.261 | 15.40 | **.002** | 0.074 |
| | | R | 34.86 | **<.001** | 0.643 | 16.00 | **<.001** | 0.443 |
| | mFus | L | 7.74 | **.001** | 0.316 | 5.91 | **.025** * | 0.185 |
| | | R | 35.70 | **<.001** | 0.612 | 9.83 | **<.001** | 0.274 |
| | pFus | L | 9.43 | **<.001** | 0.289 | 12.04 | **.005** * | 0.157 |
| | | R | 60.81 | **<.001** | 0.724 | 21.74 | **<.001** | 0.245 |
| places | COS | L | 23.39 | **<.001** * | 0.492 | 15.00 | **<.001** | 0.462 |
| | | R | 40.19 | **<.001** | 0.494 | 37.84 | **<.001** | 0.666 |
| | TOS | R | 25.27 | **<.001** * | 0.471 | 29.54 | **<.001** | 0.685 |
| bodies | ITG | L | 22.19 | **<.001** * | 0.451 | 30.64 | **<.001** | 0.754 |
| | | R | 69.82 | **<.001** * | 0.684 | 53.96 | **<.001** | 0.649 |
| | LOS | L | 27.06 | **<.001** * | 0.412 | 14.20 | **.003** * | 0.495 |
| | | R | 30.41 | **<.001** | 0.520 | 19.33 | **<.001** * | 0.183 |
| | MTG | L | 32.14 | **<.001** | 0.498 | 23.64 | **<.001** * | 0.418 |
| | | R | 41.57 | **<.001** | 0.723 | 27.63 | **<.001** | 0.474 |
| | OTS | L | 9.94 | **<.001** | 0.380 | 3.31 | .105 * | 0.133 |
| | | R | 30.06 | **<.001** | 0.593 | 8.46 | **<.001** | 0.192 |
| other objects | LOC | L | 3.70 | **.028** | 0.085 | 2.05 | .177 * | 0.035 |
| | | R | 15.11 | **<.001** | 0.402 | 2.92 | .115 * | 0.090 |

*Note.* *Greenhouse–Geisser corrected for lack of sphericity. H = hemisphere. L = Left. R = Right. Sig = Statistical significance. ROI = region of interest, for full description of ROIs see visual atlas of Rosenke et al. (2021). LOC = Lateral Occipital Complex.

**Table S11.** Summary of paired-sample t-tests examining the category-selectivity in normally-sighted individuals (the SC group) and congenital cataract reversal individuals (the CC group) of the main ROI analysis (Fig. 2).

| ROI | | H | The SC group | | | | | | | | | | | | The CC group | | | | | | | | | | | |
|---|---|---|---|---|---|---|---|---|---|---|---|---|---|---|---|---|---|---|---|---|---|---|---|---|---|---|
| | | | face vs body | | face vs object | | face vs scene | | body vs scene | | body vs object | | scene vs object | | face vs body | | face vs object | | face vs scene | | body vs scene | | body vs object | | scene vs object | |
| | | | t(7) | p | t(7) | p | t(7) | p | t(7) | p | t(7) | p | t(7) | p | t(7) | p | t(7 | p | t(7) | p | t(7) | p | t(7) | p | t(7) | p |
| faces | IOG | L | 1.67 | .114 | 1.59 | .130 | 2.95 | **.009** | | | | | | | .24 | .811 | .99 | .336 | 2.68 | **.017** | | | | | | |
| | | R | 3.08 | **.007** | 3.21 | **.005** | 4.91 | **<.001** | | | | | | | 1.75 | .099 | 2.44 | **.027** | 3.44 | **.003** | | | | | | |
| | mFus | L | 1.84 | .084 | 1.41 | .178 | 2.09 | **.053** | | | | | | | -.61 | .548 | .61 | .548 | 1.63 | .122 | | | | | | |
| | | R | 3.06 | **.007** | 3.95 | **.001** | 3.97 | **.001** | | | | | | | 1.67 | .114 | 1.67 | .114 | 2.61 | **.019** | | | | | | |
| | pFus | L | 2.92 | **.010** | .75 | .467 | 2.92 | **.010** | | | | | | | .64 | .532 | .17 | .867 | 2.61 | **.019** | | | | | | |
| | | R | 3.77 | **.002** | 5.60 | **<.001** | 5.53 | **<.001** | | | | | | | 1.65 | .118 | 1.82 | .088 | 3.36 | **.004** | | | | | | |
| places | COS | L | | | | | -3.45 | **.003** | -3.40 | **.004** | | | 2.53 | **.022** | | | | | -2.85 | **.012** | -2.55 | **.021** | | | 2.04 | .058 |
| | | R | | | | | -4.26 | **.001** | -3.75 | **.002** | | | 3.21 | **.006** | | | | | -4.31 | **.001** | -3.61 | **.002** | | | 3.12 | **.007** |
| | TOS | R | | | | | -3.24 | **.005** | -3.24 | **.005** | | | 2.94 | **.010** | | | | | -4.29 | **.001** | -3.82 | **.001** | | | 2.55 | **.021** |
| bodies | ITG | L | -3.52 | **.003** | | | | | 3.58 | **.002** | 2.85 | **.012** | | | -3.82 | **.001** | | | | | 3.97 | **.001** | 3.19 | **.006** | | |
| | | R | -4.45 | **<.001** | | | | | 5.08 | **<.001** | 4.45 | **<.001** | | | -3.36 | **.004** | | | | | 4.61 | **.000** | 3.95 | **.001** | | |
| | LOS | L | -3.53 | **.003** | | | | | 3.68 | **.002** | 3.26 | **.005** | | | -4.64 | **.000** | | | | | 3.10 | **.007** | 2.54 | **.022** | | |
| | | R | -2.91 | **.010** | | | | | 3.59 | **.002** | 3.65 | **.002** | | | -2.98 | **.009** | | | | | 3.33 | **.004** | 2.72 | **.015** | | |
| | MTG | L | -2.75 | **.014** | | | | | 5.76 | **<.001** | 3.40 | **.004** | | | -4.39 | **.000** | | | | | 3.50 | **.003** | 2.97 | **.009** | | |
| | | R | -2.39 | **.029** | | | | | 4.15 | **.001** | 3.68 | **.002** | | | -2.51 | **.023** | | | | | 3.55 | **.003** | 3.25 | **.005** | | |
| | OTS | L | -1.32 | .204 | | | | | 2.35 | **.032** | 1.67 | .115 | | | -2.68 | **.017** | | | | | 1.45 | .168 | .16 | .874 | | |
| | | R | .40 | .697 | | | | | 3.69 | **.002** | 3.47 | **.003** | | | -1.07 | .301 | | | | | 2.02 | .061 | 1.43 | .173 | | |
| other objects | LOC | L | | | -.81 | .429 | | | | | -2.12 | **.050** | -.81 | .429 | | | -.85 | .410 | | | | | -2.62 | **.018** | -.85 | .410 |
| | | R | | | 1.60 | .129 | | | | | -1.14 | .272 | -3.08 | **.007** | | | .31 | .757 | | | | | -.31 | .757 | -2.18 | **.045** |

*Note*. H = hemisphere. L = Left. R = Right. Sig = Statistical significance. ROI = region of interest, for full description of ROIs see visual atlas of Rosenke et al. (2021). LOC = Lateral Occipital Complex.

**Table S12**. Summary of two-sample t-tests comparing the response magnitude of each category between normally-sighted individuals (the SC group) and congenital cataract reversal individuals (the CC group) of the main ROI analysis (Fig. 2).

| ROI | | H | body | | | | SC vs CC group | | face | | | | SC vs CC group | | scene | | | | SC vs CC group | | other object | | | | SC vs CC group | |
|---|---|---|---|---|---|---|---|---|---|---|---|---|---|---|---|---|---|---|---|---|---|---|---|---|---|---|
| | | | SC group | | CC group | | | | SC group | | CC group | | | | SC group | | CC group | | | | SC group | | CC group | | | |
| | | | mean | SEM | mean | SEM | t(14) | p | mean | SEM | mean | SEM | t(14) | p | mean | SEM | mean | SEM | t(14) | p | mean | SEM | mean | SEM | t(14) | p |
| faces | IOG | L | 8.94 | 1.28 | 8.54 | 1.83 | 0.18 | .863 | 9.84 | 1.26 | 8.62 | 2.03 | 0.51 | .621 | 4.99 | 1.21 | 5.29 | 1.56 | -0.15 | .882 | 7.48 | 0.86 | 7.64 | 1.74 | -0.08 | .936 |
| | | R | 10.38 | 0.94 | 8.61 | 1.22 | 1.14 | .273 | 15.42 | 1.37 | 10.33 | 0.80 | 3.20 | **.008** | 4.77 | 0.66 | 4.35 | 0.80 | 0.41 | .689 | 9.86 | 1.14 | 7.20 | 0.82 | 1.90 | .080 |
| | mFus | L | 1.88 | 0.39 | 2.25 | 0.46 | -0.62 | .547 | 2.68 | 0.42 | 2.16 | 0.47 | 0.82 | .424 | 0.76 | 0.32 | 1.04 | 0.24 | -0.70 | .499 | 1.67 | 0.39 | 1.71 | 0.28 | -0.09 | .932 |
| | | R | 2.28 | 0.40 | 1.66 | 0.34 | 1.17 | .262 | 4.77 | 0.72 | 2.31 | 0.42 | 2.95 | **.013** | 0.46 | 0.31 | 0.93 | 0.25 | -1.16 | .267 | 0.97 | 0.47 | 1.38 | 0.16 | -0.81 | .438 |
| | pFus | L | 5.66 | 0.54 | 5.53 | 0.85 | 0.12 | .904 | 7.46 | 0.69 | 5.89 | 0.98 | 1.30 | .216 | 3.90 | 0.75 | 3.47 | 0.72 | 0.41 | .686 | 6.18 | 0.97 | 5.41 | 0.71 | 0.64 | .534 |
| | | R | 5.89 | 0.77 | 5.22 | 0.93 | 0.55 | .591 | 9.20 | 0.52 | 6.04 | 0.98 | 2.84 | **.016** | 2.08 | 0.61 | 2.77 | 0.53 | -0.85 | .409 | 4.12 | 0.49 | 4.90 | 0.69 | -0.93 | .371 |
| places | COS | L | 0.52 | 0.47 | 0.51 | 0.20 | 0.03 | .980 | 0.00 | 0.45 | -0.14 | 0.20 | 0.30 | .772 | 3.48 | 0.52 | 2.12 | 0.55 | 1.79 | .094 | 1.27 | 0.59 | 1.12 | 0.30 | 0.22 | .831 |
| | | R | 0.71 | 0.52 | 0.77 | 0.29 | -0.10 | .923 | 0.07 | 0.48 | 0.12 | 0.24 | -0.08 | .935 | 4.67 | 0.92 | 3.12 | 0.33 | 1.59 | .147 | 2.02 | 0.69 | 1.97 | 0.36 | 0.07 | .946 |
| | TOS | R | 1.13 | 0.86 | 1.22 | 0.31 | -0.10 | .925 | -0.30 | 1.02 | 0.11 | 0.37 | -0.38 | .711 | 5.81 | 0.94 | 5.55 | 0.58 | 0.23 | .823 | 2.12 | 0.78 | 2.97 | 0.73 | -0.81 | .434 |
| bodies | ITG | L | 21.10 | 3.87 | 13.64 | 1.25 | 1.83 | .102 | 14.26 | 3.15 | 8.53 | 0.80 | 1.76 | .116 | 4.26 | 1.92 | 3.01 | 0.51 | 0.63 | .546 | 6.78 | 1.32 | 6.71 | 0.53 | 0.05 | .964 |
| | | R | 17.83 | 1.90 | 11.70 | 1.31 | 2.65 | **.021** | 12.18 | 1.70 | 9.79 | 1.23 | 1.14 | .276 | 2.67 | 0.95 | 2.15 | 0.47 | 0.49 | .634 | 6.24 | 1.19 | 5.92 | 0.84 | 0.23 | .825 |
| | LOS | L | 16.37 | 3.01 | 9.34 | 1.17 | 2.17 | .058 | 10.37 | 2.51 | 5.93 | 1.07 | 1.63 | .136 | 3.50 | 1.60 | 3.06 | 0.46 | 0.26 | .801 | 4.77 | 1.81 | 4.53 | 0.64 | 0.12 | .904 |
| | | R | 12.12 | 2.00 | 7.15 | 1.65 | 1.92 | .076 | 7.39 | 1.34 | 6.12 | 1.70 | 0.59 | .567 | 2.14 | 0.91 | 2.18 | 1.11 | -0.03 | .977 | 3.43 | 1.17 | 4.02 | 1.59 | -0.30 | .773 |
| | MTG | L | 6.87 | 1.42 | 8.85 | 1.81 | -0.86 | .406 | 4.57 | 1.42 | 5.33 | 1.55 | -0.36 | .722 | -0.75 | 0.99 | 0.90 | 0.67 | -1.37 | .194 | 0.68 | 0.53 | 3.16 | 0.85 | -2.49 | **.029** |
| | | R | 7.89 | 1.13 | 7.99 | 1.43 | -0.05 | .958 | 5.96 | 0.73 | 6.25 | 1.12 | -0.21 | .836 | -0.43 | 0.58 | 1.09 | 0.64 | -1.77 | .099 | 1.28 | 0.61 | 3.41 | 0.84 | -2.06 | .060 |
| | OTS | L | 2.53 | 0.44 | 2.41 | 0.75 | 0.14 | .894 | 2.17 | 0.47 | 1.64 | 0.66 | 0.66 | .522 | 0.29 | 0.40 | 0.84 | 0.33 | -1.07 | .305 | 1.31 | 0.35 | 1.86 | 0.29 | -1.20 | .251 |
| | | R | 4.05 | 0.54 | 2.68 | 0.64 | 1.63 | .126 | 4.23 | 0.65 | 2.39 | 0.54 | 2.17 | **.049** | 0.66 | 0.22 | 1.05 | 0.26 | -1.13 | .279 | 1.74 | 0.38 | 1.91 | 0.39 | -0.32 | .757 |
| objects | LOC | L | 8.22 | 0.88 | 7.82 | 1.23 | 0.27 | .793 | 9.01 | 1.15 | 7.84 | 1.60 | 0.59 | .564 | 8.34 | 1.03 | 7.91 | 1.71 | 0.21 | .836 | 10.38 | 1.18 | 9.52 | 1.04 | 0.55 | .590 |
| | | R | 8.30 | 0.51 | 7.15 | 1.09 | 0.96 | .361 | 11.15 | 0.69 | 8.10 | 1.59 | 1.76 | .110 | 6.44 | 0.97 | 5.66 | 0.73 | 0.64 | .535 | 9.29 | 0.89 | 7.72 | 0.85 | 1.28 | .223 |

*Note*. Positive t-values indicate higher mean of beta-values in the SC group than in the CC group, while negative t-values indicate higher mean of beta-values in the CC group than in the SC group. H = hemisphere. L = Left. R = Right. ROI = region of interest, for full description of ROIs see visual atlas of Rosenke et al. (2021). LOC = Lateral Occipital Complex. SEM = standard error of the mean.

**Table S13**. Summary of statistical analyses (mixed-design ANOVA) for category-selectivity between normally-sighted individuals and congenital cataract reversal individuals in the ROI analysis of the **right fusiform gyrus** for different ROI sizes (Fig. S5).

| n voxels | Group | | | Category | | | | Group x Condition Interaction | | | |
|---|---|---|---|---|---|---|---|---|---|---|---|
| | F(1,14) | Sig. | Effect size (eta$^2$) | F(3,42) | Sig. | | Effect size (eta$^2$) | F(3,42) | Sig. | | Effect size (eta$^2$) |
| 30 | 1.32 | 0.270 | 0.07 | 98.87 | **<.001** | * | 0.64 | 14.35 | **<.001** | * | 0.20 |
| 40 | 0.79 | 0.390 | 0.04 | 88.85 | **<.001** | | 0.59 | 12.60 | **<.001** | | 0.17 |
| 50 | 0.62 | 0.445 | 0.03 | 85.10 | **<.001** | * | 0.57 | 11.70 | **<.001** | * | 0.16 |
| 60 | 0.40 | 0.539 | 0.02 | 79.36 | **<.001** | | 0.55 | 10.77 | **<.001** | | 0.14 |
| 70 | 0.31 | 0.586 | 0.02 | 78.20 | **<.001** | * | 0.53 | 10.24 | **<.001** | | 0.13 |
| 80 | 0.26 | 0.618 | 0.01 | 74.08 | **<.001** | | 0.51 | 9.47 | **<.001** | | 0.12 |
| 90 | 0.28 | 0.604 | 0.02 | 70.47 | **<.001** | * | 0.50 | 8.85 | **<.001** | | 0.11 |
| 100 | 0.29 | 0.602 | 0.02 | 66.66 | **<.001** | | 0.47 | 8.33 | **<.001** | | 0.10 |
| 110 | 0.29 | 0.596 | 0.02 | 63.37 | **<.001** | | 0.46 | 7.74 | **<.001** | | 0.09 |
| 120 | 0.32 | 0.583 | 0.02 | 60.24 | **<.001** | * | 0.44 | 7.36 | **<.001** | * | 0.09 |

*Note*. *Greenhouse–Geisser corrected for lack of sphericity. Sig = Statistical significance. ROI = region of interest.

**Table S14.** Summary of statistical analyses (repeated-measures ANOVA) separately in normally-sighted individuals (the SC group) and in congenital cataract reversal individuals (the CC group) in the ROI analysis of the **right fusiform gyrus** for different ROI sizes (Fig. S5).

| n voxels | The SC Group | | | | The CC Group | | | |
|---|---|---|---|---|---|---|---|---|
| | F(3,21) | Sig. | | Effect size (eta$^2$) | F(3,21) | Sig. | | Effect size (eta$^2$) |
| 30 | 52.31 | **<.001** | * | 0.57 | 52.31 | **<.001** | | 0.57 |
| 40 | 50.11 | **<.001** | | 0.53 | 50.11 | **<.001** | | 0.53 |
| 50 | 49.68 | **<.001** | | 0.52 | 49.68 | **<.001** | * | 0.52 |
| 60 | 48.07 | **<.001** | | 0.51 | 48.07 | **<.001** | | 0.51 |
| 70 | 48.39 | **<.001** | | 0.49 | 48.39 | **<.001** | | 0.49 |
| 80 | 47.35 | **<.001** | | 0.48 | 47.35 | **<.001** | | 0.48 |
| 90 | 46.26 | **<.001** | | 0.46 | 46.26 | **<.001** | | 0.46 |
| 100 | 44.78 | **<.001** | | 0.44 | 44.78 | **<.001** | | 0.44 |
| 110 | 43.73 | **<.001** | | 0.43 | 43.73 | **<.001** | * | 0.43 |
| 120 | 42.31 | **<.001** | * | 0.42 | 42.31 | **<.001** | * | 0.42 |

*Note*. *Greenhouse–Geisser corrected for lack of sphericity. Sig = Statistical significance. ROI = region of interest.

**Table S15.** Summary of paired-sample t-tests examining the category-selectivity in normally-sighted individuals (the SC group) and congenital cataract reversal individuals (the CC group) in the ROI analysis of the **right fusiform gyrus** for different ROI sizes (Fig. S5).

| n voxels | The SC group | | | | | | The CC group | | | | | |
|---|---|---|---|---|---|---|---|---|---|---|---|---|
| | face vs body | | face vs scene | | face vs object | | face vs body | | face vs scene | | face vs object | |
| | t(7) | p | t(7) | p | t(7) | p | t(7) | p | t(7) | p | t(7) | p |
| 30 | 5.65 | **.001** | 10.47 | **<.001** | 8.99 | **<.001** | 4.94 | **.002** | 6.97 | **<.001** | 5.66 | **.001** |
| 40 | 5.37 | **.001** | 9.71 | **<.001** | 8.41 | **<.001** | 4.89 | **.002** | 6.57 | **<.001** | 5.38 | **.001** |
| 50 | 5.25 | **.001** | 9.37 | **<.001** | 7.96 | **<.001** | 4.92 | **.002** | 6.70 | **<.001** | 5.24 | **.001** |
| 60 | 5.07 | **.001** | 8.95 | **<.001** | 7.51 | **<.001** | 4.92 | **.002** | 6.49 | **<.001** | 4.99 | **.002** |
| 70 | 5.01 | **.002** | 8.99 | **<.001** | 7.33 | **<.001** | 4.89 | **.002** | 6.64 | **<.001** | 4.89 | **.002** |
| 80 | 4.91 | **.002** | 8.76 | **<.001** | 7.01 | **<.001** | 4.78 | **.002** | 6.58 | **<.001** | 4.76 | **.002** |
| 90 | 4.87 | **.002** | 8.48 | **<.001** | 6.77 | **<.001** | 4.68 | **.002** | 6.57 | **<.001** | 4.60 | **.002** |
| 100 | 4.84 | **.002** | 8.27 | **<.001** | 6.49 | **<.001** | 4.54 | **.003** | 6.39 | **<.001** | 4.38 | **.003** |
| 110 | 4.75 | **.002** | 8.01 | **<.001** | 6.22 | **<.001** | 4.43 | **.003** | 6.38 | **<.001** | 4.21 | **.004** |
| 120 | 4.68 | **.002** | 7.75 | **<.001** | 5.95 | **.001** | 4.42 | **.003** | 6.38 | **<.001** | 4.11 | **.005** |

*Note*. Sig = Statistical significance. ROI = region of interest. The p-values were not corrected for multiple-comparisons.
